# Dual stop codon suppression in mammalian cells with genomically integrated genetic code expansion machinery

**DOI:** 10.1101/2023.03.26.534279

**Authors:** Birthe Meineke, Johannes Heimgärtner, Rozina Caridha, Matthias F Block, Kyle J Kimler, Maria F Pires, Michael Landreh, Simon J Elsässer

**Affiliations:** Science for Life Laboratory, Karolinska Institutet, Department of Medical Biochemistry and Biophysics, Division of Genome Biology, Stockholm 17165, Sweden; Ming Wai Lau Centre for Reparative Medicine, Stockholm node, Karolinska Institutet, Stockholm 17165, Sweden; Department of Microbiology, Tumor and Cell Biology, Science for Life Laboratory, Karolinska Institutet, Stockholm 17165, Sweden

## Abstract

Genetic code expansion via stop codon suppression is a powerful strategy to engineer proteins. Pyrrolysyine-tRNA (tRNA^Pyl^)/pyrrolysyl-tRNA synthetase (PylRS) pairs from methanogenic archaea and engineered bacterial tRNA/aminoacyl-tRNA synthetases (aaRS) pairs are used for site-specific incorporation of noncanonical amino acids (ncAAs) in response to stop codons in mammalian cells. Routinely, ncAA incorporation is achieved by transient expression of the tRNA/aaRS pair leading to heterogeneous suppression. Genomic integration of tRNA/aaRS expression cassettes for more homogenous, adjustable and reproducible levels of protein, containing one or more ncAA, will greatly benefit protein engineering, chemical control and imaging applications in mammalian cells. Here, we demonstrate that piggyBac-mediated genomic integration of archaeal tRNA^Pyl^/PylRS or bacterial tRNA/aaRS pairs, using a modular plasmid design with multi-copy tRNA arrays, allows for homogeneous and efficient, genetically encoded ncAA incorporation in diverse mammalian cell lines. We assess opportunities and limitations of using ncAAs for fluorescent labeling applications in stable cell lines. We explore simultaneous suppression of ochre and opal stop codons and finally incorporate two distinct ncAAs with mutually orthogonal click chemistries for site-specific, dual fluorophore labeling of a cell surface receptor on live mammalian cells.

## INTRODUCTION

The genetic code assigns all possible triplet codons to one of the canonical amino acids (sense codons) or to a translation stop signal (stop codons). Genetic code expansion (GCE), i.e. genetic encoding of noncanonical amino acids (ncAAs), requires reprogramming of a codon. Most commonly the amber (TAG) stop codon is repurposed, using an engineered tRNA/ aminoacyl-tRNA synthetases (aaRS) pair to deliver an ncAA-charged tRNA to the ribosome ^1,2^. The amber codon is the least abundant and most efficiently suppressed stop codon in mammals. The other two stop codons TAA (ochre) and TGA (opal), and synthetic quadruplets, have also been employed for ncAA incorporation in mammalian cells and enable dual ncAA incorporation via dual suppression ^3–9^.

In the Methanosarcinaceae family of archaea, specific amber codons in mRNAs coding for enzymes of the methanogenic pathway are suppressed by pyrrolysine tRNA (tRNA^Pyl^, encoded by the *PylT* gene)^10^. tRNA^Pyl^ and pyrrolysyl-tRNA synthetase (PylRS, encoded by the *PylS* gene) have characteristics that make them ideal for genetic code expansion in model organisms: the conserved fold of tRNA^Pyl^ allows specific interaction with PylRS independent of anticodon identity and the relatively promiscuous amino acid binding pocket of PylRS readily accommodates structurally diverse ncAAs ^11,12^. This broad substrate specificity has been further increased by protein engineering; over 100 ncAAs can be incorporated by PylRS variants (reviewed in Wan et al. ^13^). Importantly, the tRNA^Pyl^/PylRS pair is orthogonal to prokaryotic and eukaryotic tRNAs and aaRS ^14^. These features have made the pairs from *Methanosarcina mazei* (*Mma*) and *Methanosarcina bakeri* (*Mba*) widely used tools for site-specific integration of a plethora of ncAAs by amber suppression in bacterial, eukaryotic, mammalian host systems and animals ^15^.

Another approach to expand the genetic code by amber suppression relies on engineering of canonical tRNA/aaRS pairs from one organism in a host of a different clade of life. This has been achieved for both *E. coli* tyrosyl-tRNA synthetase (TyrRS) and leucyl-tRNA synthetase (LeuRS) by changing the anticodon of the cognate tRNA to CUA and sequential rounds of selection for orthogonality and efficient amber suppression. An engineered *Eco*TyrRS mutant, AzFRS, has been shown to enable effective incorporation of p-azido-phenylalanine (AzF) with either *Eco*tRNA^Tyr^_CUA_ or *B. stearothermophilus* tRNA^Tyr^_CUA_ (*Bst*tRNA^Tyr^_CUA_) ^16–19^. Further engineering of *Eco*TyrRS expanded the spectrum of ncAAs substrates ^20^. *Eco*LeuRS-derived Anap-2C (AnapRS) paired with *Eco*tRNA^Leu^_CUA_, allows introduction of the minimal fluorescent ncAA 3-(6-acetylnaphthalen-2-ylamino)-2-amino-propanoic acid (Anap) in mammalian cells ^21−^ _23._

In mammalian cell culture, components of the amber suppression machinery, i.e. the tRNA/aaRS pair and reporter or gene of interest with amber codon, are commonly expressed in transient. Lipofection methods allow easy delivery of multiple copies of the expression vectors into the cell, but the resulting populations are heterogeneous in expression levels and endogenous transcripts are outcompeted by the transgene ^24^. For prolonged and homogenous protein production, extended expression experiments, selection of homologous expression within a cell population and other applications, amber suppression cell lines with all components of the genetic code expansion machinery genomically integrated, are desirable. We have used piggyBac (PB) transgenesis to achieve genetic code expansion from stable integrants previously ^25–27^. PB transposition utilizes a deconstructed transposon cassette: cotransfection of a PB transposase (PBT) expression plasmid with a second plasmid, containing an integration cassette flanked by inverted repeats recognized by PBT, leads to random genomic integration of the cassette at TTAA sites in the host genome. Integrants are selected by antibiotic resistance markers included in the inserted cassette. While PB transgenesis is known to allow multiple integration events per cell ^28^, expression of the protein of interest is expected to be much lower and closer to high-expressing endogenous genes than expression after transient transfection. Therefore, stable genetic code expansion is commonly viewed as inefficient. Lower, genomic levels of the gene of interest may increase the competition with off-target suppression at endogenous amber codons, raising the question how selectivity towards the transgene’s stop codon can be achieved in an endogenous expression context ^29^. The comprehensive characterization of stable suppression systems for understanding the opportunities and limitations of transgenic cell lines is therefore important.

In the present study, we aimed to create a panel of representative cell lines and systematically characterize efficiency and selectivity of amber suppression using PB transgenesis. We demonstrate that stable genetic code expansion greatly varies in efficiency in different model cell lines, but can rival transient expression in specific settings and enable efficient single and dual stop codon suppression.

## RESULTS

### Integrating amber suppression machinery into the genome of human cell lines by PB transgenesis

We previously developed a plasmid system for generation of stable <u>A</u>mber <u>S</u>uppression cell lines (pAS plasmids) using *Mma* tRNA^Pyl^/PylRS (tRNA^Pyl^/PylRS hereafter) (Figure 1A) ^25,26^: One plasmid contains the *PylS* ORF for PylRS expression and a second encodes a GFP^150TAG^ reporter gene, each controlled by the EF1α promoter. Both plasmids comprise four tandem repeats of 7SK PolIII-controlled *PylT* genes for tRNA^Pyl^ expression. A puromycin or blasticidin resistance marker is expressed from an internal ribosome entry site (IRES) in the *PylS* or GFP transcripts to allow selection of integrants, respectively. The expression cassette is flanked on both ends by insulator sequences and inverted repeats for PB transposition. This plasmid design allowed for highest suppression efficiency in transient expression compared to other systems ^30^ and we have successfully employed pAS plasmids for both transient transfection and stable integration in the past ^5,8,25,27,31^.

**Figure 1.**
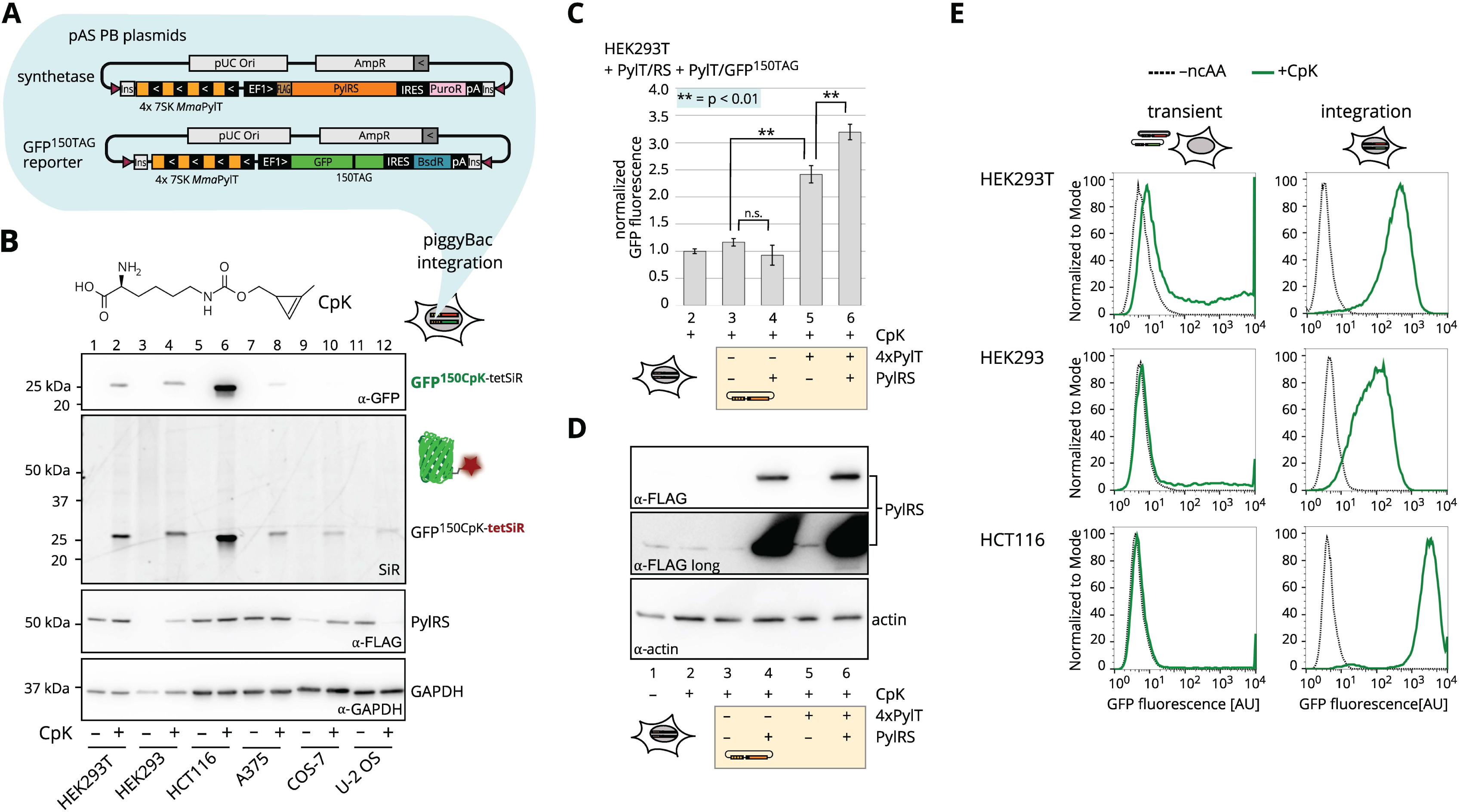
PB transgenesis integrates coding sequences for the tRNA^Pyl^/PylRS amber suppression pair into the host cell genome. A) Schematic representation of pAS (<u>A</u>mber <u>S</u>uppression) PB transgenesis plasmids. N-terminally FLAG-tagged PylRS (PylRS) is expressed under control of the EF1α promoter. An internal ribosome entry site (IRES), followed by a puromycin resistance selectable marker (PuroR), is positioned immediately downstream of PylRS. A cassette of four tandem repeats of *PylT* each controlled by h7SK PolIII promoter for tRNA^Pyl^ transcription are positioned upstream of the EF1α promoter. The entire tRNA^Pyl^/PylRS coding cassette is flanked by insulator sequences (Ins) and 3′and 5′ inverted repeats for genomic integration by PB transposase (pink triangle). The GFP^150TAG^ amber suppression reporter plasmid has the same architecture, but contains a blasticidin resistance marker (Bsd). B) Immunoblot comparing GFP^150CpK^ reporter expression in different mammalian PB integrant cell lines. The chemical structure of CpK is shown at the top of the panel. HEK293T (lanes 1 and 2), HEK293 (lanes 3 and 4), HCT116 (lanes 5 and 6), A375 (lanes 7 and 8), COS-7 (lanes 9 and 10) and U-2 OS (lanes 11 and 12) PB amber suppression integrants were cultured with 0.5 mM CpK (+, even numbered lanes) or without ncAA (–, odd numbered lanes) for 24 h. Soluble lysates were SPIEDAC labeled with 1 µM tetSiR and normalized for total protein content. SiR fluorescence was excited at 630 nm in-gel after SDS-PAGE. Immunostaining for GFP expression levels, FLAG-PylRS and GAPDH. C) Quantification of GFP fluorescence in lysates of a low selected (2 µg/ml puromycin, 500 µg/ml blasticidin), stable *PylT*, *PylS* and GFP^150TAG^ integrant, HEK293T cell line. Where indicated (yellow shading), the cell line was transiently transfected with different pAS plasmids bearing 4xPylT and/or PylRS variants. Transfections were performed in quadruplicate with 0.2 mM CpK addition for 24 h in triplicate. GFP fluorescence signal is normalized to untransfected cells cultured in presence of 0.2 mM CpK for 24 h, background fluorescence (–CpK) values were subtracted. Error bars indicate standard deviation of +CpK triplicates. Plasmids transfected by lanes: 2 no transfection, lane 3: control transfection with a pAS control plasmid, lane 4: transfection with pAS (4xPylT), lane 5: transfection with pAS_*Mma*PylRS (PylRS), lane 6: transfection with pAS_4x*Mma*PylT/RS (4xPylT and PylRS). Results from two-sided unpaired T-test are shown. D) Immunoblot for PylRS in lysates shown in C). Lanes 1 and 2: no transfection without (–) and with (+) CpK, respectively, other lanes as in C). Soluble lysates were separated by SDS-PAGE. Immunostaining for FLAG-PylRS and β-actin. E) Flow cytometry comparing amber suppression in HEK293T (top), HEK293 (middle) and HCT116 (bottom) after transient transfection (left) and PB transgenesis (right). Cells were transfected transiently for tRNA^Pyl^/PylRS and the GFP^150TAG^ reporter. Alternatively, the same plasmids were genomically integrated by PB transgenesis. GFP fluorescence from the sfGFP^150TAG^ reporter was measured after 24 h without ncAA (dashed black line) or in presence of 0.2 mM CpK (green line).

Common immortalized animal cell lines differ in their characteristics with regard to transgene expression and ease of transfection. We compare PB-mediated integration of our amber suppression system, consisting of *PylT, PylS* and the GFP^150TAG^ reporter, across a panel of commonly used cell lines. The two plasmids were cotransfected with the PBT plasmid into human embryonic kidney derived cell lines HEK293 and HEK293T, human colon cancer cell line HCT116, human melanoma A375 and osteosarcoma cell line U-2 OS as well as primate cell line COS-7. The resulting stable polyclonal pools were grown in the presence or absence of N-ε-[(2-methyl-2-cyclopropene-1-yl)-methoxy] carbonyl-L-lysine (CpK). GFP was only expressed in presence of CpK as evident from anti-GFP immunoblotting (Figure 1B, even numbered lanes). We used strain-promoted inverse electron-demand Diels-Alder cycloaddition (SPIEDAC) between CpK and tetrazine-silicon rhodamine (tetSiR) in lysates to visualize CpK incorporation ^32^. A single band corresponding to GFP^150CpK-tetSiR^ showed selective ncAA incorporation in all cell lines generated (Figure 1B). Faint additional bands indicated low level incorporation of ncAA into various endogenous proteins. The amount of GFP^150CpK^ produced greatly differed between cell lines. HCT116-derived cells produced the strongest GFP signal, followed by HEK293T and HEK293 cells. Curiously, PylRS protein levels were increased or decreased upon ncAA addition in some cell lines and PylRS expression levels did not correlate with production of full-length GFP. We did not further investigate this phenomenon. While lysates were normalized by total protein content before gel separation, GAPDH levels varied, presumably because of cell type-dependent expression patterns.

### *PylT* gene copy number is limiting in HEK293T stable amber suppression cell lines

Experiments varying copy number of tRNA and aaRS in transient transfections or viral transductions have previously helped to establish that low suppressor tRNA levels limit amber suppression efficiency ^33,34^. To determine which components are limiting amber suppression in our stable integrant system, we transfected a relatively inefficient HEK293T amber suppression cell line with different pAS constructs to increase gene copy number of *PylT*, *PylS* or both. A control pAS plasmid did not change GFP^150CpK^ or PylRS expression (lanes 1-3, Figure 1C, D). Transfection with pAS expressing PylRS lead to strongly increased PylRS protein level, but no increased amber suppression efficiency (lane 4, Figure 1C, D). Addition of pAS with either 4xPylT or 4xPylT/PylRS enhanced GFP^150CpK^ expression (lanes 5 and 6, Figure 1C, D). The GFP fluorescence increased 2.5-fold with transient transfection of pAS 4xPylT and tripled with pAS with 4xPylT/PylRS. Amber suppression did not improve by further increasing the PylT gene dosage with pAS 8xPylT, or by expressing reported optimized tRNA^Pyl^ variants M15 or A2-1 (Figure S1A) ^6,35^. Like wild-type PylRS, the chimeric *Mba/Mma*PylRS IPYE mutant had no effect on amber suppression, despite its reported higher catalytic rate ^36^ (Figure S1A). This indicates that tRNA^Pyl^ levels can limit amber suppression when they are exclusively expressed from randomly integrated pAS transgenes. The comparably high levels of amber suppression achieved in transient transfection are therefore linked to high *PylT* gene copy number leading to high levels of tRNA^Pyl^, while PylRS expression and activity are not rate-limiting.

### Comparison of transient and stable amber suppression

We further compared amber suppression efficiency of stable polyclonal HEK293, HEK293T and HCT116 cell lines with their transient transfected counterparts. After 24 h incubation with CpK, transiently transfected HEK293T cells express GFP^150CpK^ very heterogeneously: flow cytometry shows 60% GFP-positive cells, but the level of GFP fluorescence varies greatly within that population. Stable HEK293T cells produced a less extreme but more homogeneous level of GFP reporter fluorescence between 10^2^-10^3^ arbitrary units (Figure 1E). While HEK293 and HCT116 cells showed much reduced transfection efficiency, stable cell lines exhibit efficient suppression across the entire population.

We compared GFP fluorescence (indicative of the total yield of GFP) of the PB integrant cell lines to control cell lines, generated with a non-amber GFP150AAG pAS reporter plasmid, in the same HEK293T, HEK293 and HCT116 parental background (Figure S1B). The suppression efficiency, calculated as ratio of mean GFP fluorescence of GFP^150CpK^ and non-amber GFP, ranged from 11% in HEK293, to 47% and 50% in HCT116 and HEK293T cells respectively. The GFP^150CpK^ expression in stable HCT116 cells was more than 1.5times higher compared to transiently transfected HEK293T cells, suggesting that transgenic HCT116 are an excellently suited host for producing ncAA-containing proteins (Figure S1C).

We further confirmed this finding using a different ncAA and PylRS variant: The chimeric *Mma/Mba* PylRS variant AbKRS-chIPYE ^36–38^, engineered for incorporation of the photocrosslinking ncAA 3′-azibutyl-N-carbamoyl-lysine (AbK), also produced an efficiently amber-suppressing polyclonal population of HCT116 cells (Figure S1D, E).

### PB transgenesis with *E. coli* derived tRNA/aaRS amber suppression pairs in HCT116 cells

We generated pAS constructs for AzF and Anap incorporation analogous to our tRNA^Pyl^/RS two-plasmid system (Figure S2A): AzFRS was combined with *Bst*tRNA^Tyr^_CUA_ (tRNA^Tyr^_CUA_/AzFRS) and AnapRS was combined with *Eco*tRNA^Leu^_CUA_ (tRNA^Leu^_CUA_/AnapRS), both pAS were designed with a geneticin (neomycin) selectable marker. The cognate suppressor tRNA/GFP^150TAG^ reporters were also generated.

We compared amber suppression efficiency of transient transfection and PB transgenesis for AzFRS-mediated amber suppression in HCT116 cells, measuring GFP fluorescence by flow cytometry after 24 h with AzF. Transient transfection of HCT116 cells was again inefficient. Selection with geneticin and blasticidin for PB-mediated integration of tRNA^Tyr^_CUA_/AzFRS and tRNA^Tyr^_CUA_/GFP^150TAG^ produced a heterogeneous pool, but the fraction of GFP positive cells in presence of AzF increased from 4.7% in transient transfection to 39.2% (Figure 2A). Fluorescence-assisted cell sorting (FACS) of the polyclonal pool for single, strongly GFP-expressing cells, isolated clonal populations with very high amber suppression efficiency (Figure S2B). Evidently, the tRNA^Tyr^_CUA_/AzFRS pair can support extremely efficient amber suppression from integrated transgenes.

**Figure 2.**
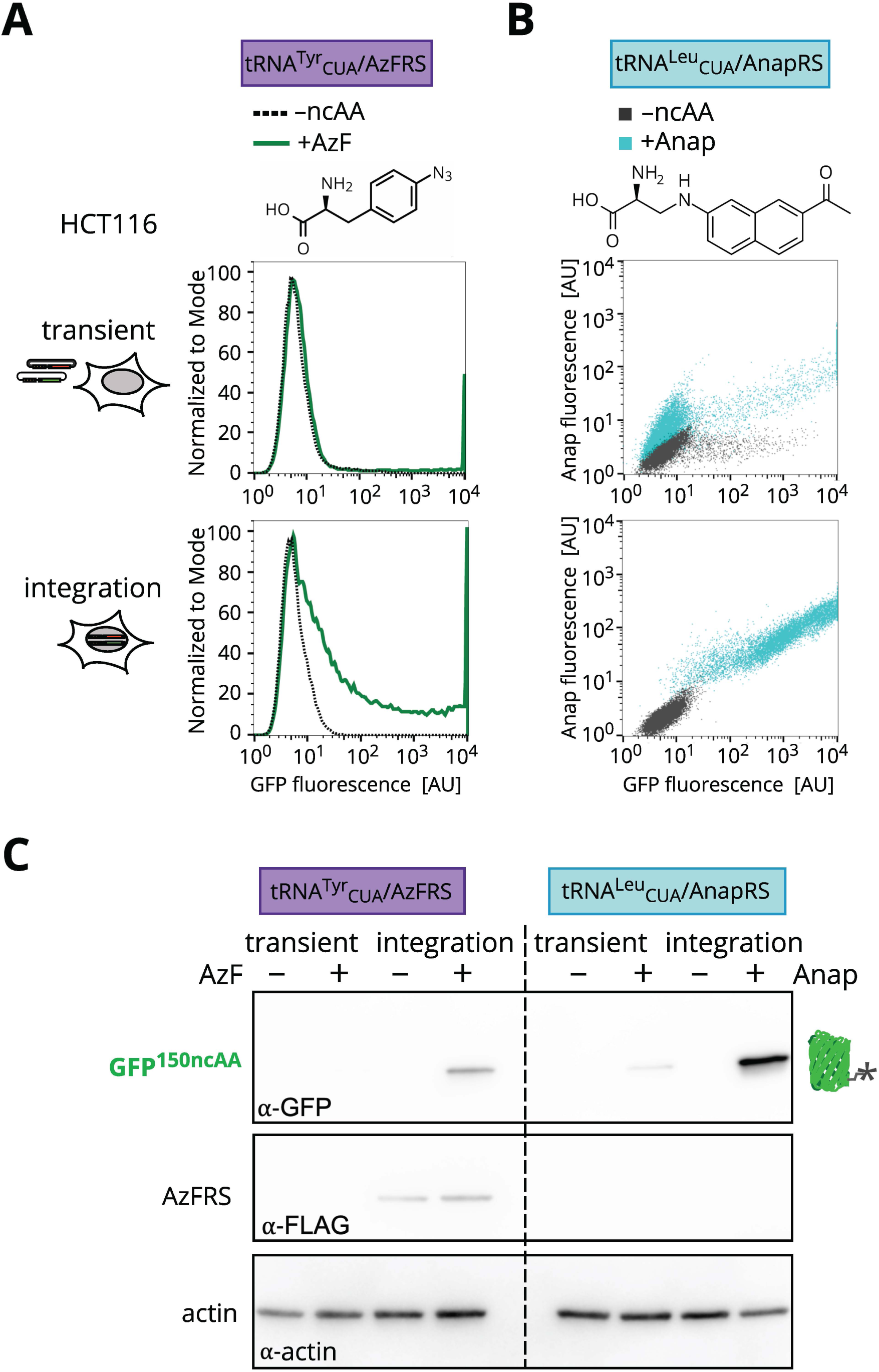
PB integration for tRNA^Tyr^_CUA_/AzFRS and tRNA^Leu^_CUA_/AnapRS mediated amber suppresion in HCT116 cells. A) Flow cytometry of amber suppression cell line generated with *E. coli* TyrRS-derived AzFRS. The chemical structure of AzF is shown. HCT116 cells were transiently transfected for tRNA^Tyr^_CUA_/AzFRS and corresponding tRNA^Tyr^_CUA_/GFP^150TAG^ reporter (transient, top). The same plasmids were integrated by PB transgenesis (integration, bottom). GFP fluorescence from the sfGFP^150TAG^ reporter was measured after 24 h without ncAA (dashed black line) or in presence of 0.5 mM AzF (green line, chemical structures shown). B) Flow cytometry of amber suppression cell line generated with *E. coli* LeuRS-derived AnapRS. The chemical structure of Anap is shown. HCT116 cells were transiently transfected for tRNA^Leu^_CUA_/AnapRS and cognate tRNA^Leu^_CUA_/GFP^150TAG^ reporter (transient, top). The same plasmids were integrated by PB transgenesis (integration, bottom). Cells were cultured with 0.05 mM Anap (cyan) or without ncAA (black) followed by a 2 h chase with ncAA-free medium prior to measurement of GFP and Anap fluorescences. C) Immunoblotting of lysates prepared from transiently transfected and PB integrant HCT116 cells cultured in presence of ncAA as in A) and B). Samples were separated by SDS-PAGE and immunostained for GFP^150ncAA^, FLAG-AzFRS and an actin loading control.

We similarly compared the polyclonal pool of stable tRNA^Leu^_CUA_/AnapRS and tRNA^Leu^_CUA_/GFP^150CpK^ integrants with transiently transfected cells. Here, we assessed both GFP and Anap fluorescence by flow cytometry (Figure 2B). The polyclonal pool of stable integrants showed strong GFP and Anap fluorescence when incubated with the ncAA for 24 h, indicating expression of GFP^150Anap^ (Figure 2B, S2C). A fluorigenic ncAA like Anap ideally allows direct detection of the protein of interest, however unincorporated and misincorporated ncAA can contribute substantially to signal intensity. To assess the contribution of misincorporated Anap in endogenous proteins, we generated a HCT116 cell line with integrated tRNA^Leu^_CUA_/AnapRS without amber suppression reporter. When cultured in presence of Anap for 24 h this AnapRS-only integrant cell line produced no GFP fluorescence, but Anap fluorescence was clearly detectable by flow cytometry despite extensive ncAA-washout before measurement (Figure S2C-E). Evidently, Anap was retained in the cell in aminoacyl-tRNA complexes or incorporated at off-target amber codons even in the absence of mRNA with a premature amber codon.

Total GFP yield, as determined by western blot, was higher in stable AzFRS and AnapRS amber suppression HCT116 cell lines, compared to transient transfection (Figure 2C). We confirmed AzFRS expression via a C-terminal FLAG tag after stable integration, while AnapRS, expressed in its originally described tagless form ^21,22^, could not be probed.

### Efficient amber-, ochre-and opal-suppressing clonal populations generated by PB transgenesis in HCT116 cells

The amber codon is most widely used in genetic code expansion in mammalian cells, because it is the least abundant stop codon (23% in the human genome) and amber suppression efficiency is superior to suppression of either of the other two stop codons, ochre (TAA) or opal (TGA) in transient transfections ^4,5,8^. We sought to explore, if genomic integration would allow for selection and isolation of efficient ochre and opal suppressor populations, a prerequisite for incorporating more than one ncAA site-specifically in response to two different stop codons in a stable cell line. We generated 4×7SK *PylT* cassettes for expression of ochre and opal suppressor tRNAs, tRNA^Pyl^_UUA_and tRNA^Pyl^_UCA_, and combined them with PylRS and their cognate GFP reporter, GFP^150TAA^ or GFP^150TGA^, respectively. In transient transfection, neither ochre nor opal suppression were efficient in HCT116 cells (Figure 3A, left column). Selection for stable integration yielded polyclonal pools of HCT116 cell lines that express GFP^150CpK^. GFP yield of the ochre and opal suppression polyclonal populations was however more than an order of magnitude lower (mean fluorescence 81 and 109 arbitrary units, respectively) than amber suppression (mean at 1687 arbitrary units) (Figure 3A, center column; Figure S3A). We isolated clones from the polyclonal population by FACS, selecting for single cells with strong GFP signal after 24 h incubation with CpK. Six amber-suppressing clones, nine ochre-and one opal-suppressing clones were recovered and compared for GFP expression by flow cytometry (Figure S3A, B). For each clone, addition of CpK resulted in homogenous GFP fluorescence at higher intensity than observed for the polyclonal pool. The most efficiently suppressing clones for each stop codon are shown in Figure 3A (right column). Despite the clonal selection, maximal ochre and opal suppression remained roughly one order of magnitude lower than amber suppression (Figure S3B). This confirmed the observation from transient transfection experiments, that ochre and opal codons suppression is less efficient ^3-^ _9._

**Figure 3.**
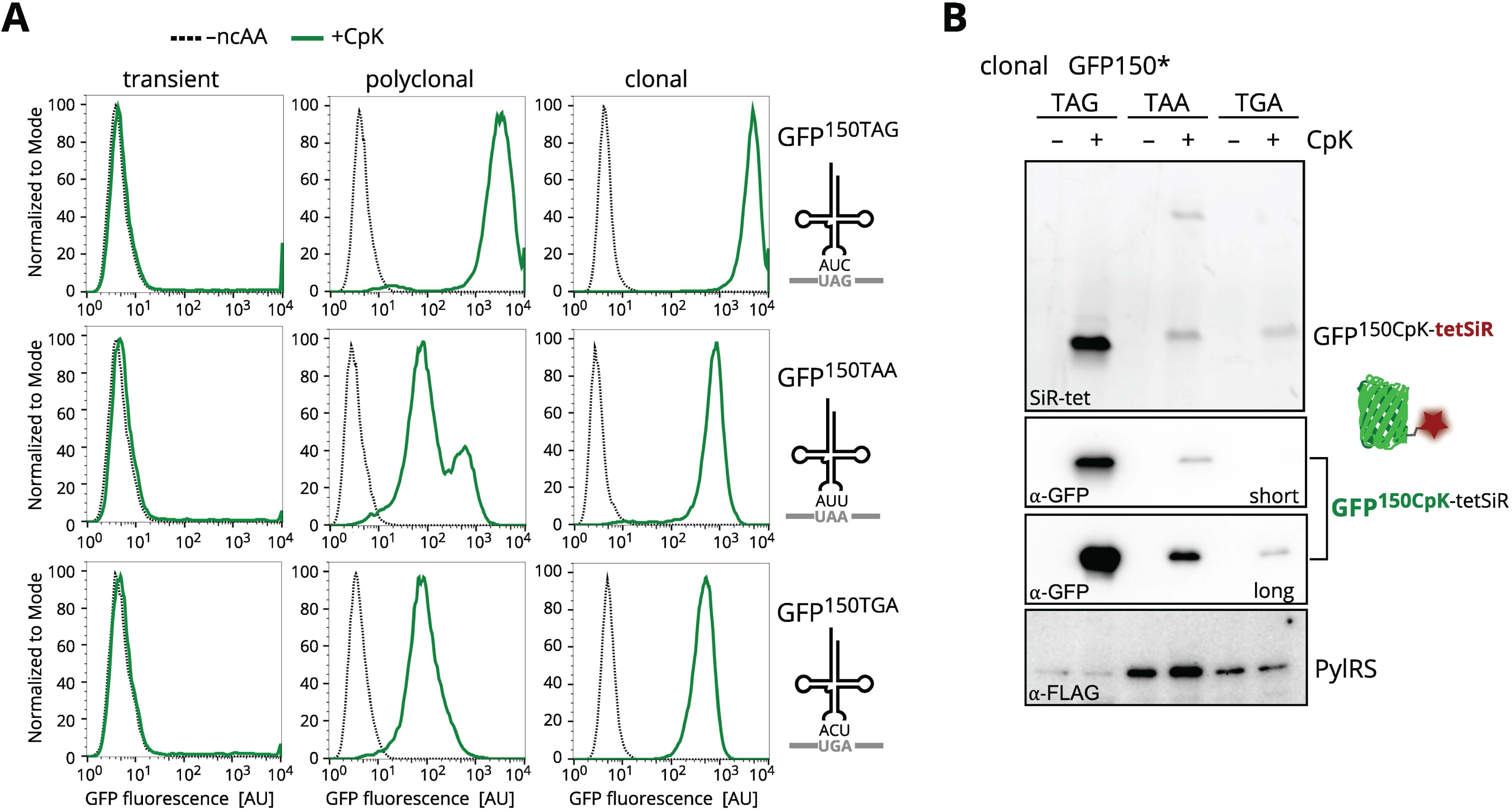
Generation of efficient ochre and opal suppressor clonal HCT116 populations by PB transgenesis. A) Flow cytometry comparing amber (TAG, top), ochre (TAA, middle) and opal (TGA, bottom) suppression in HCT116 cells after transient transfection (left) and PB transgenesis (middle and right). GFP fluorescence was measured without ncAA (dashed black line) or 24 h after addition of 0.2 mM CpK (green line). Left: Transient transfection with pAS plasmids for tRNA^Pyl^, PylRS and the GFP^150stop^ reporter, suppressor tRNAs have the indicated CUA, UUA or UCA anticodons to match the UAG, UAA and UGA stop codons in the respective GFP reporter mRNA (grey). Center: using the same pAS plasmids, expression cassettes producing PylRS, tRNA^Pyl^_CUA_, tRNA^Pyl^_UUA_ or tRNA^Pyl^_UCA_ (black) and the corresponding GFP150stop reporter, were genomically integrated by PB transgenesis. Right: Clonal populations, isolated from the polyclonal pools (middle) by FACS, after 24 h incubation with 0.2 mM CpK. Best clones are shown for amber, ochre and opal suppression, respectively. The other recovered clones are compared in Figure S3. Data for amber suppression after transient transfection and stable integration in HCT116 cells are reproduced from Figure 1E. B) Immunoblot of lysates from clonal stop codon suppressor HCT116-derived cell lines shown in (A). The asterisk (*) indicates a stop codon. Clonal populations were cultured with 0.5 mM CpK for 72 h. GFP^150CpK^ was SPIEDAC labeled with 1 µM tetSiR in lysate followed by SDS-PAGE and immunostaining for GFP expression and FLAG-aaRS expression.

Efficient suppression of the target stop codon is typically considered the most important variable for optimization, but the selectivity for ncAA labeling applications also crucially depends on low misincorporation at endogenous stop codons ^29^. In order to compare the on-versus off-target incorporation of ncAA, we compared the best clones for each stop codon after extended CpK incorporation. Cell lysates were labeled by SPIEDAC with tetSiR during lysis (Figure 3B). The most prominent band corresponds to GFP^150CpK-tetSiR^ in each of the three stop codon-suppressing cell lines. The ochre suppression clonal cell lysate also featured a distinct band, suggesting a particularly strongly labeled endogenous protein. It is interesting to note, that endogenous opal codons do not appear to be more efficiently suppressed by tRNA^Pyl^ than the other two stop codons, despite the opal codon being known to show highest leakiness, i.e., readthrough with endogenous tRNAs ^33,39,40^. In summary, productive ochre and opal suppression is possible to achieve in stable settings. Yet, as expected, amber suppression of the reporter GFP is markedly more efficient while exhibiting similarly low levels of incorporation into endogenous proteins.

### Dual suppression clonal cell line for amber and ochre co-suppression

In recent years, several approaches have successfully implemented dual suppression of two different stop codons in transiently transfected mammalian cells, allowing for site-specific incorporation of two distinct ncAAs ^3–8^. Encouraged by selection of ochre and opal suppressor cell lines expressing high levels of GFP^150CpK^, we decided to investigate if dual suppression can be implemented in a stable setting. The compatibility of tRNA^Tyr^_CUA_/AzFRS and tRNA^Leu^_CUA_/AnapRS pairs engineered from *E. coli* in combinations with archaeal tRNA^Pyl^/PylRS has been explored in detail using transient transfection ^4^. Choosing the two most efficiently suppressed stop codons (Figure 3A), we explored the combination of the ochre suppressor tRNA^Pyl^_UUA_/PylRS pair with a compatible orthogonal amber suppressor tRNA/aaRS pair for dual integration. We integrated tRNA^Pyl^_UUA_/PylRS, tRNA^Tyr^_CUA_/AzFRS and a GFP^102TAG150TAA^ dual suppression reporter into HCT116 cells (Figure 4A). After co-selection with puromycin, geneticin and blasticidin, a few cells showed increased GFP fluorescence in the polyclonal pool after 48 h with CpK and AzF. We isolated single cells with high GFP fluorescence by FACS and recovered clonal populations with a varied range of dual suppression efficiency (Figure S4A). The most efficient clone showed robust, CpK-and AzF-dependent expression of GFP^102AzF150CpK^ (Figure 4B, C and S4B, C). No cross-incorporation of ncAA was observed; there are no GFP-positive cells after incubation with only one of the ncAAs (Figure 4B, C). We purified GFP expressed in presence of AzF and CpK from the clonal cell line (yield 80 ng/ml culture) and confirmed incorporation of both ncAAs by intact mass spectrometry (Figure 4D). In addition to CpK, GFP^102AzF150CpK^ contains AzF, which can also be used for bioorthogonal labeling. The azido group reacts with alkynes in Cu(I) catalyzed azide-alkyne cycloaddition (CuAAC) or with strained alkynes in copper independent strain promoted azide-alkyne cycloaddition (SPAAC). We visualized purified GFP^102AzF150CpK^ by SPIEDAC and SPAAC with tetSiR and dibenzylcyclooctyne-PEG4-5/6-tetramethyl-rhodamine (DBCO-TAMRA), respectively (Figure 4E). In summary, stable cell lines are useful for production of dual suppressed proteins.

**Figure 4.**
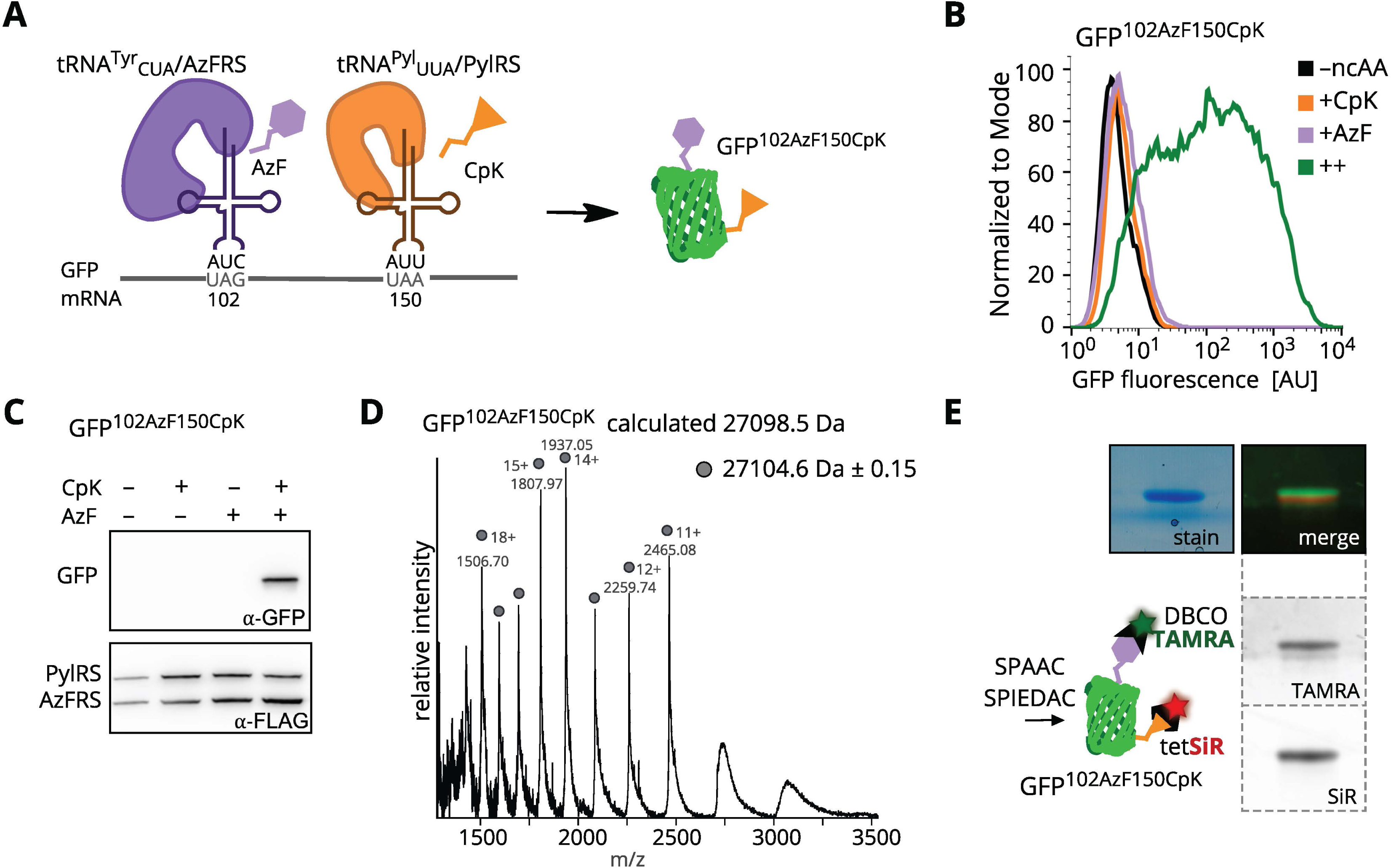
Dual suppression clonal cell line for GFP^102AzF150CpK^ expression in HCT116 cells. A) Scheme illustrating amber and ochre co-suppression in HCT116 cells with tRNA^Pyl^_UUA_/PylRS ochre suppressor pair, tRNA^Tyr^_CUA_/AzFRS amber suppressor pair and a GFP^102TAG150TAA^ reporter using pASs integrated by PB transgenesis. Addition of AzF and CpK allows GFP^102AzF150CpK^ production. B) Flow cytometry of the HCT116 PtRNA^Pyl^_UUA_/PylRS, tRNA^Tyr^_CUA_/AzFRS and GFP^102TAG150TAA^ clone F6 grown without ncAA (–ncAA, black), with 0.2 mM CpK (+CpK, orange), with 0.5 mM AzF (+AzF, violet) or both ncAAs (++, green) for 48 h. C) Immunoblot for lysates prepared from the GFP^102TAG150TAA^ integrant clone F6 grown without ncAA, with 0.2 mM CpK, with 0.5 mM AzF or both ncAAs for 48 h. Lysates were separated by SDS-PAGE followed by immunostaining for GFP expression and FLAG-aaRS expression. D) Intact mass determination for purified GFP^102AzF150CpK^ expressed in the dual suppressor clonal cells. Calculated theoretical mass and determined mass are indicated. E) GFP^102AzF150CpK^ purified from the PB dual suppressor HCT116 clone by SPAAC and SPIEDAC with 1 µM DBCO-TAMRA and 1 µM tetSiR, respectively. In-gel fluorescence was excited at 520 nm (for TAMRA) and 630 nm (for SiR) after separation by SDS-PAGE, fluorescence merge and coomassie-stained gel are shown.

### Amber suppression cell lines for cell surface receptor expression

We have so far described PB cell line generation with an easily detectable, well expressed and stable GFP reporter. To demonstrate the robustness of our approach, we chose a range of other proteins involved in different cellular processes. We generated stable HCT116 amber suppression cells lines for expression of a number of intracellular proteins of interest, either with premature TAG or STELLA tag (N-terminal TAG) and C-terminal HA-tag ^31^: The cytoskeletal protein tubulin, heterochromatin associated DAXX, the histone variant CENP-A and the microproteins PIGBOS and SARS2-CoV-M (Figure S5A). We used the PylRS “AF” variant in these cell lines to incorporate axial trans-cyclooct-2-ene-L-lysine (TCO*K, Figure 5A) ^41,42^. Polyclonal integrant populations were selected and all proteins were detectable by immunoblotting for their HA-tag, with expected variation in expression levels. SPIEDAC labeling with tetSiR or tetrazine-AF488 (tet488) in cell lysates shows specific bands for the TCO*K-labeled proteins according to their respective size, albeit endogenous proteins appear to be labeled with similar or even higher intensity (Figure S5B, C). Fluorescence microscopy of fixed and permeabilized cells labeled with methyl-tetrazine-BDP-FL (metetBDPFL) and counterstained for HA further showed that bioorthogonal labeling created a strong ambient background signal across cytosol and nucleus while a specific signal from the amber suppressed protein was only marginally brighter. This is most evident for the Golgi-restricted CoV-M protein (Figure S5D) ^31^. Therefore, the strong expression and selective suppression of our GFP reporter must be considered an outlier compared to many biologically relevant proteins that are desirable to label using amber suppression. Evidently, the known large plasmid copy number, ten to hundred thousand copies per cell ^43^, after transient transfection provides a much larger window for selective suppression of the target amber codon than expression from genomic PB integrants. Nevertheless, we considered that mislabeling of endogenous C-terminal amber stop codons may generate predominantly intracellular background and using a cell-impermeable dye may allow selective labeling of target proteins on the cell surface.

**Figure 5.**
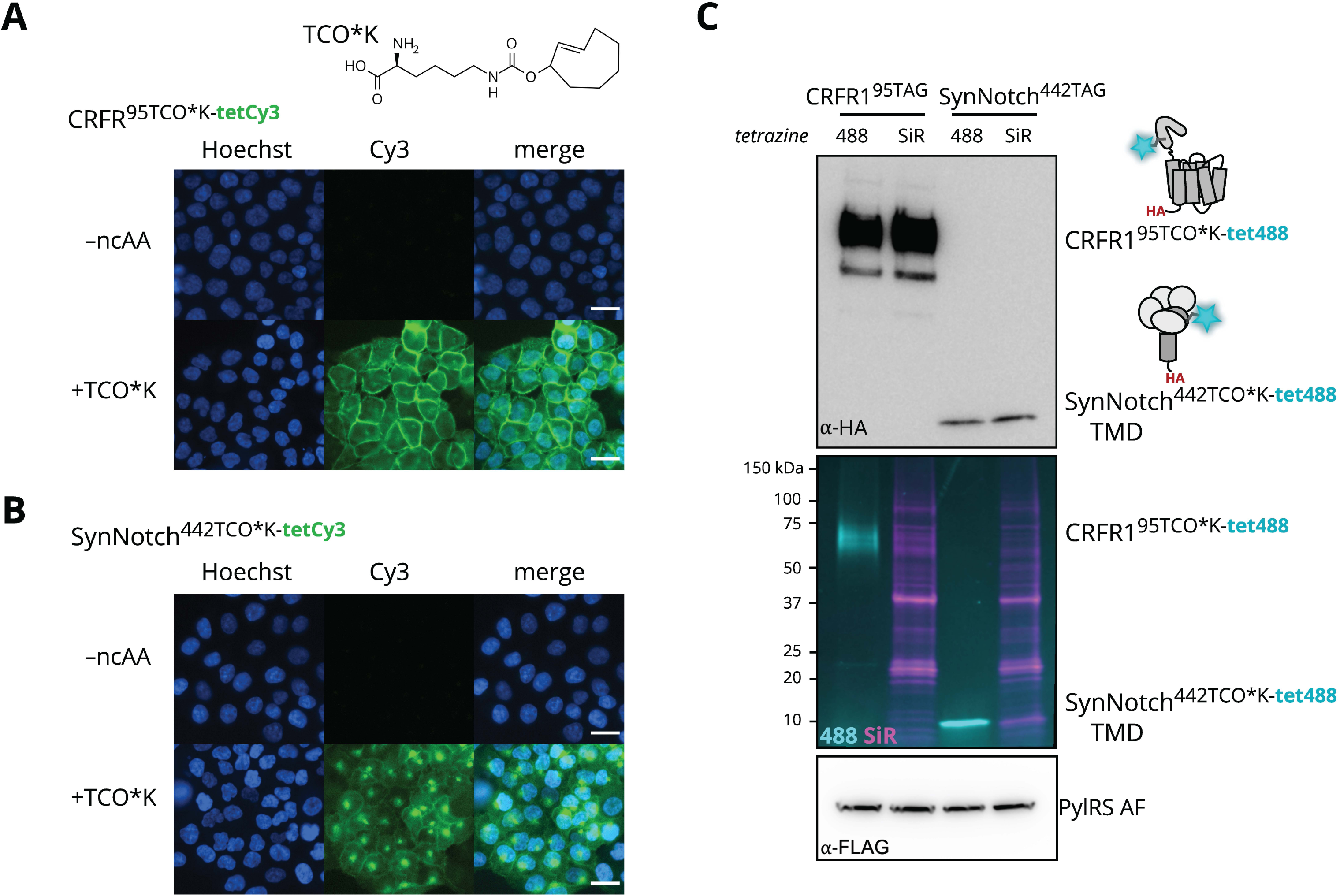
Amber suppression cell lines for bioorthogonal labeling of cell surface receptors. Fluorescence microscopy imaging of HCT116 PB integrant cell lines for cell surface receptor expression by amber suppression with (A) tRNA^Pyl^/PylRS AF and CRFR1^95TAG^ or (B) tRNA^Pyl^/PylRS AF and SynNotch^442TAG^. Integrant populations recovered after selection were cultured with 0.1 mM TCO*K for 48 h, chased in ncAA free medium for 2 h prior to SPIEDAC labeling with 1 μM tetCy3. Cells were fixed in PFA and nuclei were counterstained with Hoechst 33342 prior to imaging on a Nikon Ti2. White scale bars indicate 25 μm. C) Immunoblot for the same cell lines as in A) and B) expressing CRFR1^95TCO*K^ or SynNotch^442TCO*K^. Cells were cultured with 0.1 mM TCO*K for 48 h. The cell surface receptors were either labeled by SPIEDAC with 2 μM membrane impermeable tet488 before lysis or with 1 μM tet-SiR in cell lysate. Cleared lysate aliquots were separated by SDS-PAGE and tet488 surface fluorescence and tetSiR lysate labeling visualized with 460 nm (for 488, cyan) and 630 nm (for SiR, magenta) excitation, respectively. Immunostaining for the C-terminal HA-tags confirms specific SPIEDAC surface labeling of CRFR1^95TCO*K^ and SynNotch^N442TCO*K^. Immunostaining for FLAG-PylRS AF is shown. The SynNotch illustration was simplified by omitting the N-terminal LaG17 fusion.

To test this hypothesis, we used two cell surface receptors for generation of stable amber suppression HCT116 cell lines: First, the class B GPCR corticotropin-releasing factor type 1 receptor (CRFR1). CRFR1 A95TAG has previously been used in amber suppression studies for ncAA-mediated bioorthogonal labeling on the surface of live cells ^6,8,44,45^.

Second, a minimal synthetic Notch receptor (SynNotch): An amber codon at N442 in our construct (N1713 in human *Notch1*) places the ncAA on the cell surface close to the membrane spanning helix of the C-terminal fragment generated by proteolysis during maturation of the receptor ^8,46^.

Polyclonal stable cell lines for each receptor were cultured with TCO*K for 48 h and live cells were SPIEDAC-labeled with tetrazine-Cy3 (tetCy3) and subsequently fixed for microscopy. CRFR1^95TCO*K^-and SynNotch^442TCO*K^-expressing cells showed a strong, TCO*K-dependent tetCy3 signal on the surface of cells (Figure 5A, B). Expression levels were much more homogeneous across the population compared to transient transfection ^8^. The same CRFR1^95TAG^ and SynNotch^442TAG^ cell lines were labeled live with tet488 and lysates were labeled with tetSiR for comparison (Figure 5C). While the tet488 signal was highly specific for the cell surface receptor and overlapped with the HA-immunostaining signal, tetSiR in lysates preferentially labeled endogenous proteins. In summary, stable amber suppression cell lines provide limited opportunity for selective bioorthogonal labeling of intracellular proteins, but enable efficient suppression and labeling of extracellular receptors, offering an ideal platform for biophysical studies of membrane receptors in their native environment.

### Dual site-specific ncAA incorporation and dual color labeling by combined amber and ochre suppression in transgenic HCT116 cells

Building on our promising results for combining amber and ochre suppression with tRNA^Pyl^_UUA_/RS and tRNA^Tyr^_CUA_/AzFRS for the GFP reporter, we attempted generation of stable dual suppression cell lines in HCT116 cells for SynNotch. The SynNotch^204TAA442TAG^ amber and ochre double mutant can incorporate two distinct ncAAs for dual bioorthogonal derivatization when transiently cotransfected with appropriate, orthogonal tRNA/aaRS pairs. The incorporation sites are designed to place the two ncAAs in close spatial proximity to each other, but on separate parts of the mature, proteolytically processed SynNotch receptor ^8^.

Initial attempts to generate PB transgenic cell lines for SynNotch^204TAA442TAG^-HA failed, even when we exchanged the ochre suppressor tRNA^Pyl^_UUA_ for the M15 mutant ^6^, which we had found to enhance ochre suppression efficiency before ^8^. We recovered PylRS and AzFRS expressing populations after selection, yet were unable to detect ncAAs or HA-tag (data not shown). To facilitate selection of integrants with SynNotch receptor expression by fluorescence-activated cell sorting (FACS), we exchanged the C-terminal HA-tag for a GFP-tag. We transfected HCT116 cells with M15_UUA_/PylRS, tRNA^Tyr^_CUA_/AzFRS, M15_UUA_/SynNotch^204TAA442TAG^-GFP and PBT. After selection with geneticin, puromycin and blasticidin, GFP fluorescence could be detected in a small number of cells after culturing in presence of AzF and CpK for 3 days. From this population, we isolated single cells by FACS for highest GFP fluorescence (Figure S6A). The best clone recovered showed increased GFP fluorescence with both ncAAs over a 12 to 72 h time course (Figure S6B). Flow cytometry and imaging of this clone confirmed that GFP fluorescence was not the result of cross-incorporation, readthrough with endogenous amino acids or a secondary translation start, as it localized to the cell membrane and depended on addition of both CpK and AzF (Figure 6A, S6C).

**Figure 6.**
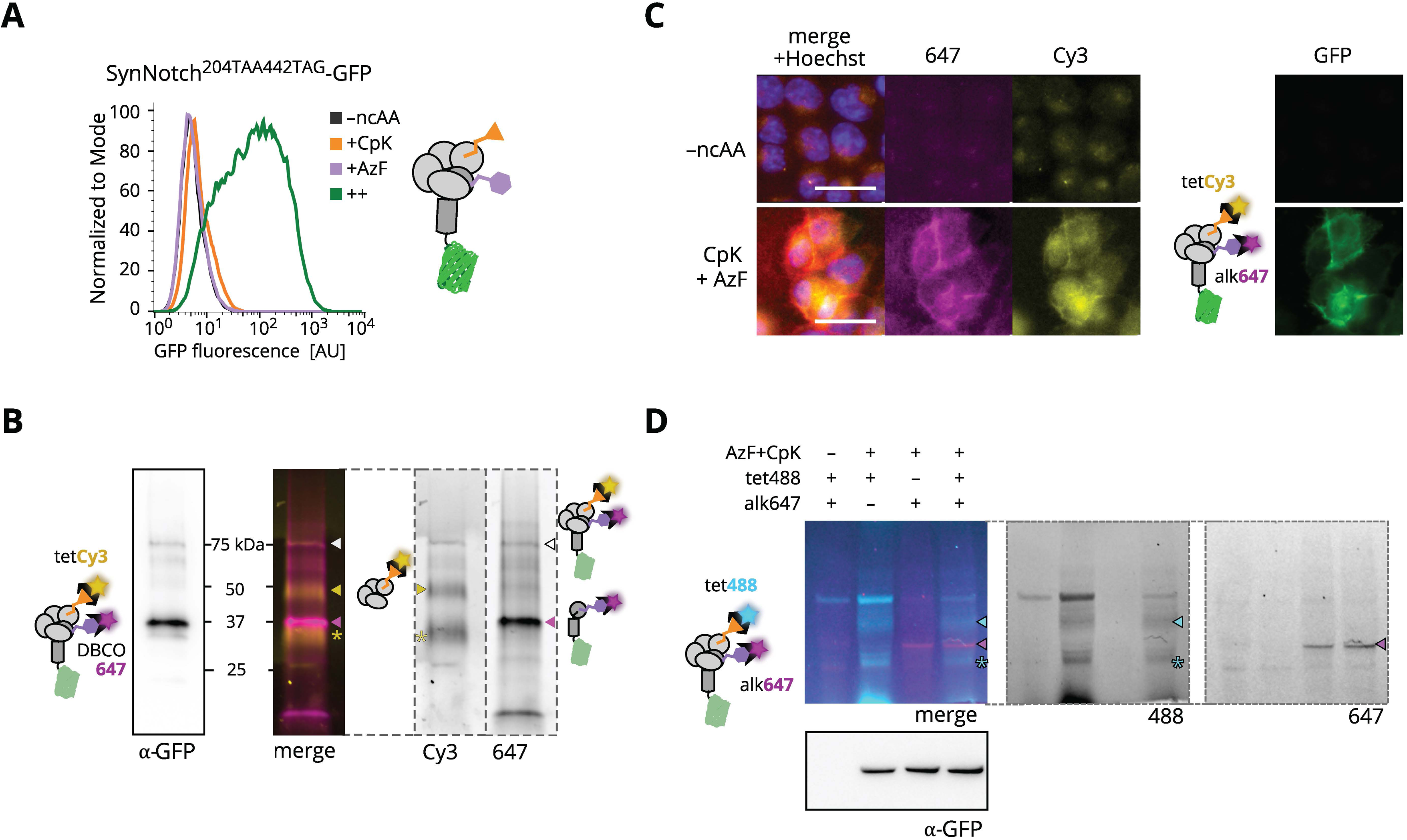
Dual suppression HCT116 clonal cell line for SynNotch^204CpK442AzF^ expression and live-cell dual bioorthogonal labeling. A) The HCT116 dual suppressor clone, with M15_UUA_/PylRS ochre suppressor pair, tRNA^Tyr^_CUA_/AzFRS amber suppressor pair and SynNotch^204TAA442TAG^-GFP integrated via PB transgenesis, was cultured 72 h without ncAA (–ncAA, black), with 0.2 mM CpK (+CpK, orange), 0.5 mM AzF (+AzF, purple) or both ncAAs (++, green) and analyzed by flow cytometry for GFP fluorescence. The positions of both ncAAs in the mature receptor are illustrated in the schematic. SynNotch illustrations are simplified by omitting the N-terminal LaG17 fusion. B) SynNotch^204CpK442AzF^-GFP purified from the dual suppression HCT116 integrant clone. Bioorthogonal labeling by SPAAC and SPIEDAC with 2 μM DBCO-647 and 2 μM tetCy3, respectively. In-gel fluorescence of the bead-captured protein after separation by SDS-PAGE, 520 nm (for Cy3) and 630 nm (for AF647) excitation and merged signals are shown, as well as immunoblotting against the C-terminal GFP. The different domains and processing states of SynNotch are indicated - white arrow: 81 kDa pre-processed SynNotch^204CpK442AzF^; yellow arrow: 43 kDa N-terminal domain with 204CpK labeled by tetCy3; magenta arrow: 38 kDa C-terminal TMD-GFP fusion with 442AzF labeled by DBCO647; yellow asterisk: additional SynNotch N-terminal fragment after proteolytic cleavage of LaG17 ^8^. C) Fluorescence microscopy for dual bioorthogonal labeling of SynNotch^204CpK442AzF^-GFP expressed in the clonal dual suppressor clone. Cells were cultured with 0.2 mM CpK and 0.5 mM AzF for 72 h, labeled with 2 μM tetCy3 by SPIEDAC and 2 μM alk647 by CuAAC, respectively. Prior to imaging on a Nikon Ti2, nuclei were counterstained with Hoechst 33342. White scale bar indicates 25 μm. See also Figure S6E. D) SDS-PAGE gel separation of lysates from SynNotch^204CpK442AzF^-GFP expressing dual suppressor HCT116 clone. SPIEDAC and CuAAC labeling with 2 µM tet488 and 2 µM alk647 on live cells cultured with 0.2 mM CpK and 0.5 mM AzF for 72 h. In-gel fluorescence after separation, excitation 460 nm (for AF488) and 630 nm (for AF647) and immunoblotting against the C-terminal GFP are shown. Arrows indicate the different processing states of SynNotch: Cyan arrow indicates the 43 kDa N-terminal domain containing 204CpK-tet488 after SPIEDAC, magenta arrow indicates 38 kDa C-terminal TMD-GFP fusion comprising 442AzF-alk647, asterisk indicates an additional SynNotch N-terminal fragment without LaG17 ^8^.

We validated the functionality of the two ncAAs for dual bioorthogonal labeling with tetCy3 and DBCO-AF647 (DBCO-647) by SPIEDAC and SPAAC, respectively, on SynNotch^204CpK442AzF^ - GFP purified from the clone (Figure 6B). In SDS-PAGE the two proteolytic fragments of SynNotch are separated. The Cy3 and 647 fluorescence signals are isolated which demonstrates that each receptor domain is selectively labeled and confirms site-specific incorporation of the two ncAAs. As expected, the DBCO647 signal from the AzF labeled C-terminal transmembrane domain overlapped with anti-GFP immunostaining signal. We observe two diffuse bands for the glycosylated N-terminal domain, due to partial proteolytic cleavage after the nanobody domain^8^. A small proportion of unprocessed SynNotch^204CpK442AzF^-GFP was also captured, visible as a faint high molecular weight band in both fluorescence channels and immunoblotting. We validated the band assignment by dual suppression of transiently transfected SynNotch^204TAA442TAG^-GFP and SPIEDAC labeling on live cells, which resulted in the same pattern (Figure S6D).

### Live-cell dual color bioorthogonal labeling of a receptor in stable cell lines

Next, we investigated, if the clonal stable dual suppression cell line produced sufficient SynNotch^204CpK442AzF^-GFP for live cell labeling. Cells were grown in presence of both ncAAs for 72 h, labeled with tetCy3 and alkyne-AF647 (alk647) and imaged for fluorescence of both dyes (Figure 6C). We detected overlapping, ncAA-dependent cell surface signals for all three fluorophores associated with SynNotch^204CpK442AzF^-GFP. The Cy3, 647 and GFP fluorescence signals were dependent on addition of both ncAA. When CuAAC was performed on live cells, both GFP and Cy3 signals were decreased; highlighting the potential of copper to damage cellular proteins even at low concentration (Figure S6E). Using a copper-free SPAAC reaction with DBCO-467 as an alternative for labeling AzF, however, was not successful, as it led to strong unspecific labeling of endogenous proteins (Figure S6F) ^46^.

SynNotch^204CpK442AzF^-GFP expressing cells labeled by SPIEDAC and CuAAC under live conditions yielded the same band pattern observed for affinity purified SynNotch^204CpK442AzF^-GFP when separated by SDS-PAGE (Figure 6D). Two ncAA-dependent bands with tet488-fluorescence are visible for the N-terminal SynNotch domain, due to partial proteolysis as previously observed. An additional higher molecular weight band represented an artifact of tet488 surface labeling since it was present in all dye-treated samples, even in absence of ncAAs (Figure 6D). After CuAAC labeling a single fluorescent band is visible, corresponding to C-terminal TMD-GFP, comprising N442AzF specifically labeled by alk647 (Figure 6D). SPAAC labeling of SynNotch^204CpK442AzF^-GFP with DBCO-647 lead to a diffuse signal across the entire lane, owing to the unspecific DBCO reactivity we also observed during microscopy (Figure S6G) ^46^. Taken together, our results demonstrate that dual suppression and bioorthogonal derivatization of two distinct ncAAs with two mutually orthogonal reactions is possible in a stable cell line with integrated suppression machinery. The resulting fluorescence signal strength was limited by the low efficiency of dual stop codon suppression and by the limited choice of mutually orthogonal reactions.

## DISCUSSION

Systematic evaluation of suppression systems and mammalian host cell lines enabled us to generate stable cell lines with expanded genetic codes for suppression of all three stop codons using PB transgenesis. We established a universal two-plasmid design for generating mammalian cell lines with the choice of three distinct orthogonal tRNA/Synthetase pairs. This versatility should enable incorporation of virtually any genetically encoded ncAA described to date in stable cell lines. While stop codon suppression, especially for ochre and opal codons, can be initially low after drug selection, highly efficient clones can be isolated. In the present study, we did not systematically assess DNA copy number and resulting relative expression levels of PylRS, tRNA^Pyl^ and the target protein in a larger number of clones. Consequently, a universal recipe for the optimal dosage of tRNA, aaRS and gene of interest cannot be derived from our data, and the best condition may be highly context specific with respect to cell line, target protein, and ncAA used. Nevertheless, we find that maximizing tRNA and gene of interest expression are determinants of ncAA incorporation efficiency and selectivity. In addition, position and context of the stop codon have been shown to greatly influence suppression efficiency and hence are important parameters to optimize ^29^.

Our approach benefits from the efficiency of PB transgenesis. Optimal clones can be isolated by selection without a priori knowledge of the ideal copy number configuration. As an alternative to genomic integration, an episomal self-replicating plasmid has been used to long-term express the amber suppression machinery in human hematopoietic stem cells ^47^. This approach, however, will likely require continuous selection to maintain stable transgene copy number and efficient amber suppression. Stable integration and clonal selection, including by PB transposition, has been also used to generate amber suppression CHO cell lines efficiently expressing ncAA-modified antibodies for drug conjugation ^48,49^.

Building on the toolbox created in this study, we proceeded to demonstrate site-specific incorporation and chemical labeling of two distinct ncAAs in a stable mammalian cell line. Our study highlights opportunities and limitations of dual suppression systems, requiring two selective, efficient and mutual orthogonal incorporation machineries. The relatively lower efficiency of ochre and opal stop codon suppression limits overall yield of dual suppressed protein ^4,5,8,9^. We chose AzF for encoding the CuAAC handle because of its efficient incorporation (Figure 2A, S2B), but this likely limited the fluorescent labeling efficiency on live cells. We have previously shown that a genetically encoded picolyl-azide provide superior CuAAC reactivity with alkyne dyes ^45^, but orthogonal machineries to introduce picolyl-azide and TCO*K or CpK do not exist to date.

It is an important goal of the field to achieve selective labeling of amber codons in target proteins expressed at endogenous level or even endogenously edited genes. Our experiments with the GFP reporter demonstrate that, under most optimal conditions, endogenously expressed proteins can be ncAA-labeled with high efficiency and good selectivity. However, our data also underscore that such an ideal scenario is unlikely to apply to the average cellular protein. Our methodology provides most attractive applications for labeling proteins on the plasma membrane. Site-specifically incorporated ncAAs on the cell surface face little competition from misincorporated ncAAs at endogenous stop codons due to the fact that C-termini of plasma membrane proteins are mostly intracellular or posttranslationally processed. Our stable genetic code expansion technology, together with fluorescent bioorthogonal labeling on the cell surface, provides a valuable toolbox to investigate the localization, movement and clustering of membrane receptors on the cell surface. Dual-color labeling will enable new applications for studying conformational dynamics and multimeric assemblies of cell surface proteins.

## AUTHOR CONTRIBUTIONS

BM and SJE conceived and designed experiments. BM, JH and MFP generated plasmids. BM, JH and RC generated stable cell lines. JH and BM performed fluorescence microscopy. BM, RC, KJK and MFB per. BM and SJE analyzed data. ML performed intact mass spectrometry measurements and analyzed data. BM and SJE analyzed data. BM prepared figures. BM, JH and SJE wrote the manuscript. All authors read the manuscript.

## ACKNOWLEDGEMENTS

We thank all members of the Elsässer lab for input and the other groups in the Division of Genome Biology for their support: the J. Bartek lab for access to a Tecan Infinite M200 Pro Plate Reader and a Nikon Eclipse Ti2 Microscope, and the O. Fernandez-Capetillo lab for access to a GE AI600 gel-imager. Research was funded by Vetenskapsrådet, Sweden (2015– 04815); the Ming Wai Lau Center for Reparative Medicine, Sweden; the Ragnar Söderbergs Stiftelse, Sweden; Stiftelsen för Strategisk Forskning (FFL7); and the Knut och Alice Wallenbergs Stiftelse, Sweden (2017–0276).

## MATERIAL AND METHODS

### Materials availability

A list of PiggyBac integrant cell lines has been deposited to Mendeley data. The DOI is listed in the key resources table. Clonal populations isolated in this study are listed in the key resources table and available on request. Most plasmids used in this study have been deposited to Addgene, all others are available on request.

### Cell Culture Material

HEK293 cells (human, sex unknown, epithelial-like embryonic kidney), HEK293T cells (human, sex unknown, epithelial-like embryonic kidney), HCT116 cells (human, male, colon cancer), A375 cells (human, female, malignant melanoma), U-2 OS cells (human, female, osteosarcoma) and COS-7 cells (*Cercopithecus aethiops*, sex unknown, fibroblast kidney) were maintained in Dulbecco’s™ modified Eagle’s medium (DMEM, GlutaMAX, Thermo) supplemented with 10 % (v/v) FBS at 37 °C and 5 % CO_2_ atmosphere.

## Methods

### DNA constructs

The *M. mazei* tRNA^Pyl^/PylRS expression and super folder GFP^150TAG^ (referred to as GFP^150TAG^ and GFP throughout) reporter constructs with four tandem h7SK driven *PylT* repeats, coding for U25C tRNA^Pyl^, have been described previously ^5^. Analogous constructs for amber suppression by AnapRS and AzFRS with the respective cognate tRNA^Leu^_CUA_ or tRNA^Tyr^_CUA_ were generated for this study. The plasmids share a common architecture and are here collectively referred to as “pAS” (<u>A</u>mber<u>S</u>uppression) plasmids: The aaRS, reporter or gene of interest coding sequences are controlled by EF1α promoter and followed by a internal ribosome entry site (IRES) that allows expression of a downstream selection marker. A cassette with four tandem repeats of the tRNA gene, controlled by the human 7SK PolIII promoter, are placed upstream of the EF1αpromoter in anti-sense orientation. A schematic of this architecture is shown in Figure 1A. All DNA constructs were verified by Sanger sequencing. Refer to the key resources table for plasmids used and generated in this study and their Addgene accession number.

### Cell Culture and Transfection

HEK293, HEK293T, HCT116, A375, COS-7 and U-2 OS cells were maintained in Dulbecco’s modified Eagle’s medium (DMEM, GlutaMAX, Thermo) supplemented with 10 % (v/v) FBS (Sigma) at 37 °C and 5% CO_2_ atmosphere. For transfection (1.5-2.0) x 10^5^ cells/ml were seeded 24 h before transient transfection with 1 µg plasmid DNA/ml using TransIT-LT1 (Mirus) according to the manufacturer’s instructions. In transient transfection experiments, ncAAs were added at the time of transfection and cells were harvested after 24, 48 or 72 h, as indicated.

### Generation of PB-mediated stable integration cell lines

Cell lines with PiggyBac (PB) integrated transgenes were generated as described ^25,26^: Parental cells were seeded one day prior to transfection and transfected with desired pAS plasmids and PiggyBac transposase (PBT) in a 4:1 ratio. Earliest 48 h after transfection, cells were split and selected with different concentrations of appropriate antibiotics for 7 days (ranges: 2-5 μg/ml puromycin (VWR), 500-4000 μg/ml blasticidin (Invivogen), 2000-3000 μg/ml geneticin (G418, Sigma)). Stable integrant cell populations were recovered from the highest submissible selection condition in DMEM + 10% FBS (v/v). Polyclonal populations were cultured with ncAA for 24, 48 or 72 h, as indicated, and characterized by flow cytometry for reporter fluorescence and immunoblotting for reporter or protein of interest and PylRS. A list of cell lines generated in this study can be found on Mendeley data (https://doi.org/10.17632/tgc7mbv5xp).

### Noncanonical amino acids

Working stocks of ncAAs were prepared in 100 mM NaOH with 15% DMSO (v/v) to 100 mM. Anap working stock was prepared to 20 mM in the same buffer. Chemical structures of N-ε-[(2-methyl-2-cyclopropene-1-yl)-methoxy] carbonyl-L-lysine (CpK), p-azido-phenylalanine (AzF), 3-(6-acetylnaphthalen-2-ylamino)-2-amino-propanoic acid (Anap) and axial trans-cyclooct-2-ene-L-lysine (TCO*K) are shown in Figures 1, 2 and 5, respectively. Refer to the Key Resource Table for supplier and CAS numbers.

### SPIEDAC lysate labeling, SDS-PAGE and immunoblotting

Aliquots of cell lysate were separated on 4-20 % Tris-glycine gels (BioRad) and transferred to nitrocellulose membranes (BioRad). Cells were lysed in RIPA (for intracellular proteins) or PBS with 0.1% triton X-100 (v/v) (cell membrane proteins) supplemented with 1x cOmplete proteinase inhibitor (Roche). The insoluble fraction was removed by centrifugation. To compare amber suppression of PB transgenic integrants generated from different human cells, the soluble fraction of whole cell lysates were normalized for total protein content (BCA assay, Pierce). For SPIEDAC labeling in lysate, 1 μM tetrazine-Silicon rhodamine (tetSiR) (Spirochrome) was added to the lysis buffer. Lysates cleared by centrifugation were mixed with 6x Laemmli buffer and denatured for 10 min at 37°C, or at 95°C to completely denature GFP. After separation, gels were exposed at 460 nm, 520 nm and 630 nm in a GE AI600 imager for in-gel fluorescence. Expression of GFP reporter, HA-or GFP-fusion proteins and FLAG-aaRS was confirmed by immunoblotting with antibodies against GFP (Santa Cruz, sc-9996), HA-HRP (Roche, 12013819001), FLAG-HRP (Sigma, A8592), β-actin (cell signaling, 4970), GAPDH (Millipore, AB2302) and corresponding secondary HRP-conjugated antibodies when needed (BioRad). Annotated, preprocessed immunoblotting data can be found on Mendeley data (https://doi.org/10.17632/tgc7mbv5xp).

### Quantification of GFP expression

HEK293T-derived cells, selected for pAS_4x*Mma*PylT/RS and pAS_4x*Mma*PylT/GFP^150TAG^ PB transposition (low antibiotic concentration: 2 µg/ml puromycin, 500 µg/ml blasticidin), were transiently transfected with an additional pAS plasmid in quadruplicate. Transfected cells were grown in absence or presence (triplicate) of 0.2 mM CpK for 24 h. Cells were lysed in RIPA buffer with 1x cOmplete protease inhibitor (Roche), the insoluble fraction was removed by centrifugation. GFP bottom fluorescence of an aliquot was measured in Tecan Infinite M200 pro plate reader (excitation 485 nm, emission 518 nm). Fluorescence measurements were normalized to total protein content of each sample as determined by Pierce BCA assay kit (Fisher Scientific) on the same sample. Average GFP fluorescence was calculated from the technical triplicates, the GFP fluorescence measured in absence of CpK was subtracted as background fluorescence and values were normalized to average GFP fluorescence measured for the untransfected amber suppression cell line.

### Flow cytometry and fluorescence-assisted cell sorting (FACS)

Cells were trypsinized after the indicated time of culturing in presence of ncAAs and resuspended in PBS + 10% FBS (v/v). Flow cytometry was carried out on a Beckmann NAVIOS flow cytometer and analyzed in FlowJo^TM^ software (version 10.6.2) (BD Life Science). Polyclonal populations selected with antibiotics after PB transgenesis were cultured with the cognate ncAAs for 24 h (GFP^150TAG^, GFP^150TAA^, GFP^150TGA^), 48 h (GFP^102TAG150TAA^) or 72 h (SynNotch^204TAA442TAG^-GFP). Single cells were isolated by gating for top 0.4-5% GFP fluorescence signal on a SONY SH800 cell sorter. Clonal populations were expanded in DMEM + 10% FBS and analyzed by flow cytometry and immunoblotting for GFP and aaRS expression.

### Intact mass spectrometry

The dual suppression GFP^102TAG150TAA^ clonal integrant cell line was cultured in the presence of 0.2 mM and 0.5 mM AzF for 96 h. Cells were lysed in RIPA buffer supplemented with 1x cOmplete protease inhibitor (Roche). The insoluble fraction was removed by centrifugation. Expressed GFP was captured on GFP-Trap_MA magnetic beads (ChromoTEK), washed with RIPA buffer, PBS + 500 mM NaCl and PBS prior to elution in 1 % (v/v) acetic acid.

Purified GFP^102AzF150CpK^ samples were directly infused into a Waters Synapt G1 traveling-wave IM mass spectrometer (MS Vision, Almere, The Netherlands), with an m/z range of 500 to 4000 Th. Mass spectra were recorded in positive ionization mode. The capillary voltage was maintained at 1,5 kV and the sample cone was 100 V. The extraction cone voltage was 4 V. The source temperature was maintained +30°C. The trap and transfer collision energies were 10 V. The trap gas was argon at a flow rate of 4 mL/h. Data was analyzed with MassLynx version 4.1 (Waters).

### Bioorthogonal labeling on beads

GFP^102AzF150CpK^ and SynNotch^204CpK442AzF^-GFP were captured on GFP-Trap_MA magnetic beads (ChromoTEK) from cell lysate of the respective PB stable clonal cell lines cultured in presence of 0.5 mM AzF and 0.2 mM CpK for 72 h. Cells were lysed and the insoluble fraction was removed by centrifugation. Beads were washed with RIPA buffer, PBS + 500 mM NaCl and PBS. SPIEDAC and SPAAC were carried out on bead-bound proteins simultaneously by adding 1 μM tetSiR (spirochrome) or 1 μM tetCy3 (Click Chemistry Tools) and 1 μM DBCO-TAMRA (Jena Bioscience) or 1 μM DBCO-647 (Jena Bioscience) for 10 min on ice. Excess dye was washed off and bead-bound proteins were eluted with 1 % (v/v) acetic acid for SDS-PAGE separation. Equal amounts of GFP purified from different samples were separated on 4-20% Tris-glycine gels (BioRad) and exposed for in-gel fluorescence at 520 nm and 630 nm in a GE AI600 imager. The gel was stained with InstantBlue (Expedeon) to visualize GFP^102AzF150CpK^ bands or transferred to a nitrocellulose membrane for immunostaining against GFP to detect SynNotch^204CpK442AzF^-GFP.

### Live-cell surface SPIEDAC and CuAAC labeling for in-gel fluorescence and fluorescence microscopy

Surface SPIEDAC labeling was performed with 1-2 µM tet488 (Jena Bioscience) or 1-2μM tetCy3 (Click Chemistry Tools) in DMEM + 10% FBS (v/v) for 30 min at 37 °C. Where indicated SPAAC was carried out with 2 μM DBCO-AF647 (DBCO-647, Click Chemistry Tools) in DMEM +10% FBS (v/v) for 30 min at 37 °C. For subsequent CuAAC in the dual suppression cell line, cells were washed with PBS and labeling was performed with 50 µM CuSO_4_, 250 µM THPTA, 2.5 mM sodium ascorbate and 2 µM alk647 (Jena Bioscience) for 10 min at 22 °C. Cells were washed and collected in PBS and lysed in PBS with 0.2% triton X-100 (v/v) and 1x complete protease inhibitor (Roche) on ice. Lysates cleared by centrifugation were mixed with 6x Laemmli buffer and denatured for 10 min at 37°C, or at 95°C to completely denature GFP. Proteins were separated by SDS-PAGE and in-gel fluorescence measured as described above. Alternatively, for cells grown on poly-L-lysine coated 18-well glass imaging slides (Ibidi), cells were washed with PBS, fixed in paraformaldehyde (PFA) 4% (v/v) and counterstained with 2 µM hoechst 33342 prior to imaging in PBS on a Nikon Eclipse Ti2 inverted widefield microscope, using a 20x (0.75 NA) objective and filter sets for DAPI, GFP, Cy3 and Cy5 fluorescence. Images were processed in Fiji ^50^. To ensure comparability of fluorescence signal between images from different samples, images were acquired at constant settings for each filter set after initial adjustment. Images collected with each filter set were processed as stacks in Fiji.

## SUPPLEMENTARY MATERIAL AND METHODS

### Noncanonical amino acids

*N6*-[[2-(3-Methyl-3*H*-diazirin-3-yl)ethoxy]carbonyl]-L-lysine (AbK, CAS: 1253643-88-7, SiChem, SC-8034) was prepared as described for the other ncAAs.

### Intracellular SPIEDAC labeling and immunofluorescence microscopy

HCT116 PB integrant polyclonal population expressing tRNA^Pyl^, PylRS AF and gene of interest (GOI) were seeded at 15 000 cells per well on poly-L-lysine coated 96 well imaging plates (Ibidi) in presence of 0.1 mM TCO*K for 48 h. Subsequently, the cells were washed with PBS, fixed in 4% PFA) for 10 min, permeabilized in PBS with 0.1% triton X-100 (v/v) (PBS-T), blocked in 2% BSA in PBS-T and stained with 0.5 μM methyltetrazine-BDP-FL (metetBDPFL) (Jena Bioscience) for 1 h at RT, incubated with primary anti-HA antibody (Santa Cruz, sc-7392) followed by incubation with secondary 555-coupled antibody (Life Technologies, A-31570) and 2 µM Hoechst 33342 (Life Technologies). The cells were washed and imaged in PBS on a Nikon Eclipse Ti2 inverted widefield microscope, using a 20x (0.75 NA) objective and filters sets for DAPI, AF488, Cy3 and Cy5 fluorescence.

**Figure S1.**
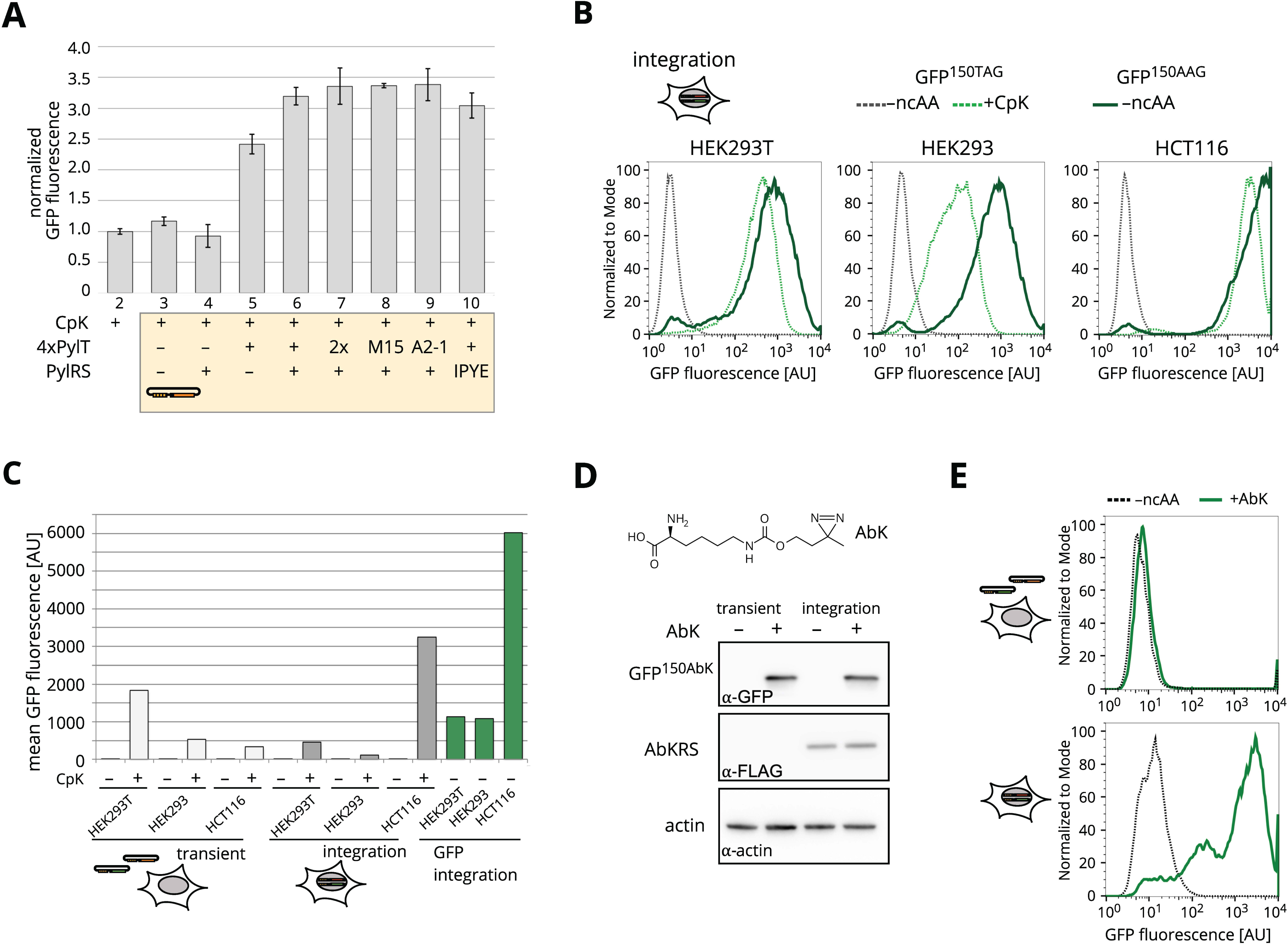
PylRS is not limiting for amber suppression PB stable cell lines and PB integration of AbKRS. Related to Figure 1. A. Quantification of GFP fluorescence in lysates of low selected (2 µg/ml puromycin, 500 µg/ml blasticidin) stable integrant HEK293T cell line. Where indicated (yellow shading), the cell line was transiently transfected with different pAS plasmids bearing *PylT* for tRNA^Pyl^ and/or *PylS* for different PylRS variants. For each condition, quadruplicate transfections were performed. For three of the four samples medium was supplemented with 0.2 mM CpK, all samples were harvested 24 h post transfection. GFP fluorescence signal is normalized to that of untransfected cells cultured in presence of 0.2 mM CpK for 24 h, background fluorescence (–CpK) values were subtracted. Error bars indicate standard deviation. Plasmids transfected by lanes: 1 no transfection, 2 control plasmid GFP^102TAA133TGA150TAG^, 3 PylRS (Addgene #154762), 4 4xPylT (Addgene #140008), 5 4xPylT/RS (Addgene #140009), 6 8xPylT/RS (Addgene #140008), 7 4xM15T/PylRS (Addgene #174890), 8 4xA2-1T/PylRS (A2-1 ^1^), 9 4xPylT/IPYE_*Mba/Mma*chPylRS ^2^. B. Flow cytometry comparing amber suppression in HEK293T (left), HEK293 (middle) and HCT116 (right) to the same cell type after PB transgenesis with a GFP150K reporter (Addgene #193313, AAG coding for lysine in position 150). Plasmids were integrated into the genome by PB transgenesis. Reporter GFP fluorescence was measured after 24 h without ncAA (dashed grey line) or in presence of 0.2 mM CpK (dashed light green line) or for non-amber GFP150K without ncAA (dark green line). C. Quantification of mean GFP fluorescence in flow cytometry samples shown in Figure 1D and PB integration control cell lines with GFP150K (Addgene #193313) shown in Figure S1B generated in HEK293T, HEK293 and HCT116 cells. D. Immunoblotting of amber suppression with PylRS variant AbKRS-chIPYE (AbKRS). The chemical structure of *N6*-[[2-(3-Methyl-3*H*-diazirin-3-yl)ethoxy]carbonyl]-L-lysine (AbK) is shown. HCT116 cells transiently transfected with PylT/AbKRS and PylT/GFP^150TAG^ plasmids (transient, left), or polyclonal stable integrants with the same plasmids (integration, right) were grown without ncAA (–AbK) or presence of 0.5 mM AbK for 24 h. Immunostaining for GFP expression levels, FLAG-AbKRS and β-actin. E. Flow cytometry of HCT116 cells transiently transfected with PylT/AbKRS and PylT/GFP^150TAG^ plasmids (transient, top), or polyclonal stable integrants with the same plasmids (integration, bottom) were grown in absence of ncAA (black) or in presence of 0.5 mM AbK (green) for 24 h.

**Figure S2.**
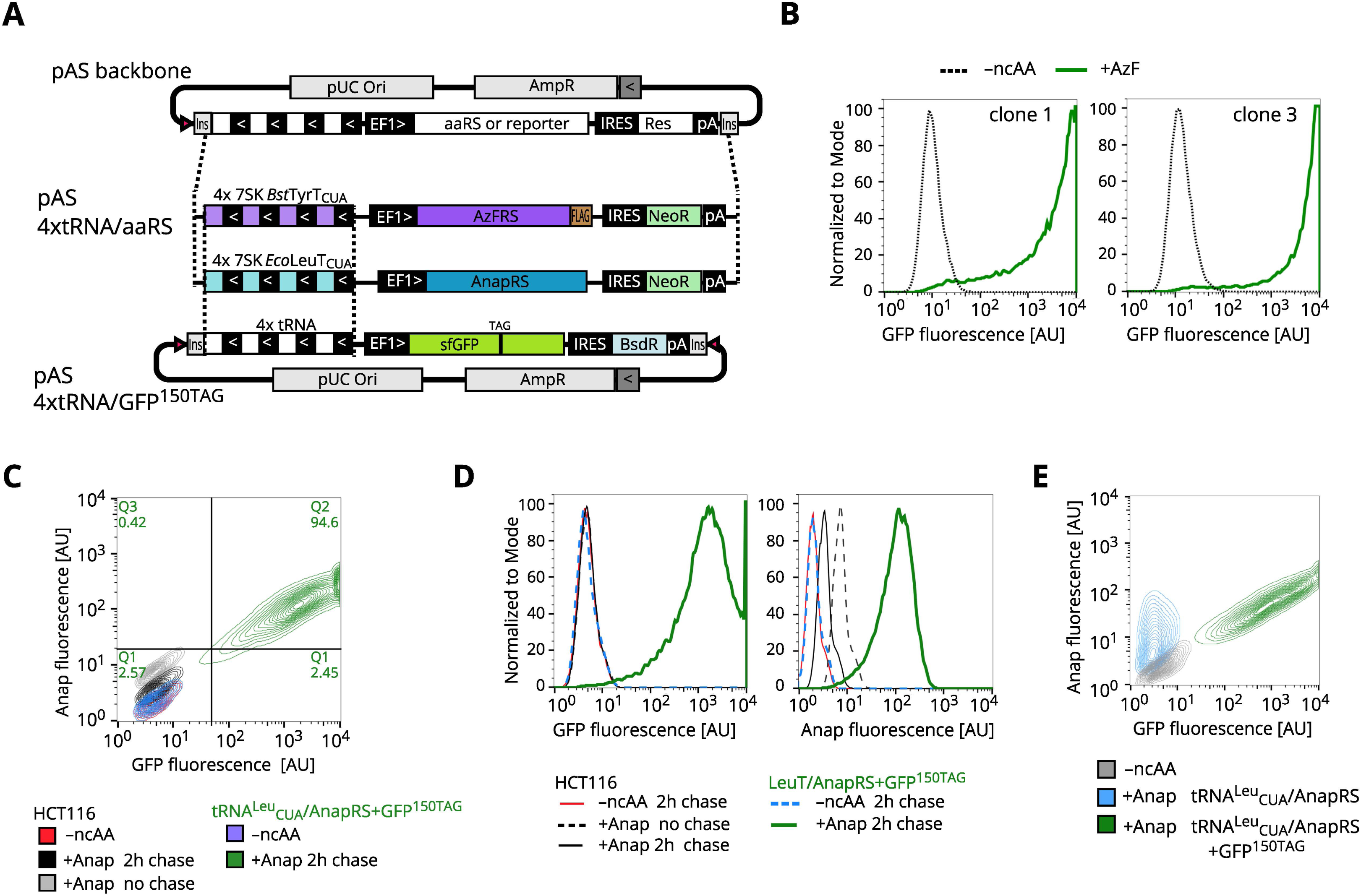
AzFRS PB integrant amber suppression clones and limitations of AnapRS PB integration in HCT116 cells. Related to Figure 2. A. Schematic representation of pAS constructs for AzF and Anap incorporation. AzFRS or AnapRS are expressed under EF1α promoter control. An IRES (internal ribosome entry site) followed by a neomycin resistance (NeoR) is positioned immediately downstream of the aaRS. A cassette of four tandem copies of orthogonal tRNA (*B. stearothermophilus* TyrT_CUA_ (coding for tRNA^Tyr^_CUA_) or *E. coli* LeuT_CUA_ (coding for tRNA^Leu^_CUA_) for 7SK PolIII driven expression is integrated upstream of the EF1α promoter. We chose to add external 7SK PolIII promoters leading each tRNA, even for *Bst* tRNA^Tyr^_CUA_ which contains an internal B-box for PolIII transcription ^3^. The entire 4xtRNA/aaRS coding cassette is flanked by insulator sequences (Ins) and 3’ and 5’ inverted repeats for genomic integration by PBT (pink triangle). Plasmids for GFP^150TAG^ amber suppression reporter with either tRNA^Tyr^_CUA_ or tRNA^Leu^_CUA_ expression cassettes have the same architecture, but contain a blasticidin resistance gene (Bsd). B. Flow cytometry of amber suppression clonal cell lines isolated from the AzFRS, cognate tRNA^Tyr^_CUA_ and GFP^150TAG^ reporter integrant polyclonal population shown in Figure 2. GFP fluorescence from the GFP^150TAG^ reporter was measured in the two clonal cell lines after 24 h without ncAA (dashed black line) or in presence of 0.5 mM AzF (green line). C. Flow cytometry of amber suppression cell line generated with AnapRS, cognate tRNA^Leu^_CUA_ tRNA and GFP^150TAG^ reporter compared to the parental HCT116 cells. Cells were grown in presence of 0.05 mM Anap for 24 h and either measured directly or after a 2 h chase with ncAA-free medium. The 2D plot for GFP and Anap fluorescence shows the wild type HCT116 populations without Anap (-ncAA, red), with Anap (+Anap, grey) and with Anap washed out (+Anap, chase, black). The tRNA^Leu^_CUA_/AnapRS+GFP^150TAG^ integrant stable cell line is shown in blue (-ncAA) and green (+Anap). D. Same data as in C) presented as histogram plots separating GFP and Anap fluorescences. E. Flow cytometry of amber suppression cell line generated with AnapRS, cognate tRNA^Leu^_CUA_ and a GFP^150TAG^ reporter (green) compared to an integrant cell line with tRNA^Leu^_CUA_/AnapRS only (no reporter, blue). Cells were grown in presence of Anap for 24 h and measured after a 4 h chase with ncAA-free medium. The “no reporter” cell line retains Anap.

**Figure S3.**
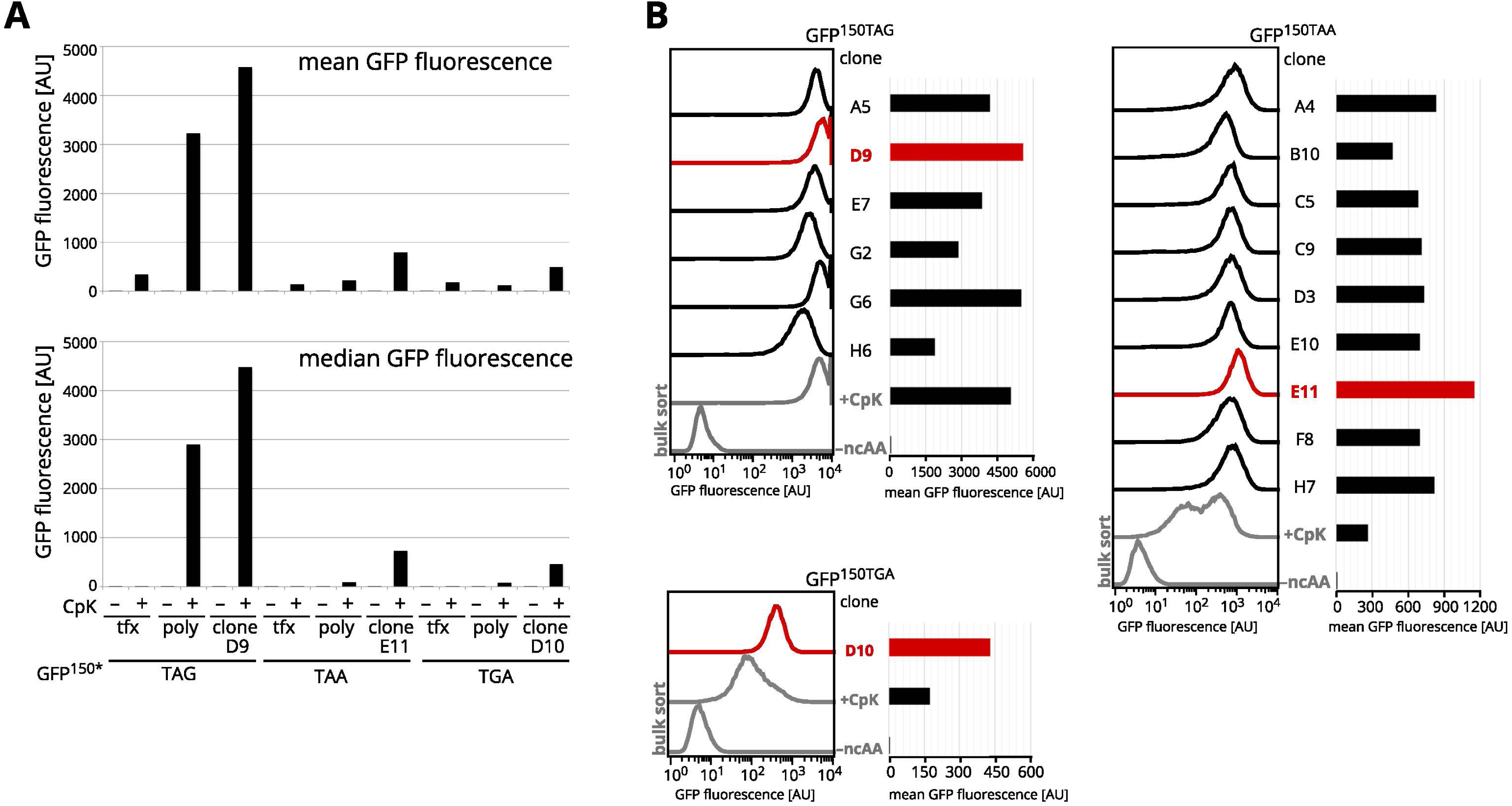
Identification of best amber, ochre and opal suppressor clonal HCT116 populations. Related to Figure 3. A. Quantification of mean and median GFP fluorescence measured in Figure 3A. B. Comparison of GFP fluorescence generated by amber, ochre or opal suppression in the dual suppression clones sorted from polyclonal pools of tRNA^Pyl^_TXX_/PylRS and GFP^150^* reporter HCT116 integrants generated by PB transgenesis. The clonal populations were grown in presence of 0.2 mM CpK for 24 h. The clones used in Figure 3A are highlighted in red. For comparison a population grown from 100 cells is also shown in grey, as –ncAA control and approximation of the polyclonal pool.

**Figure S4.**
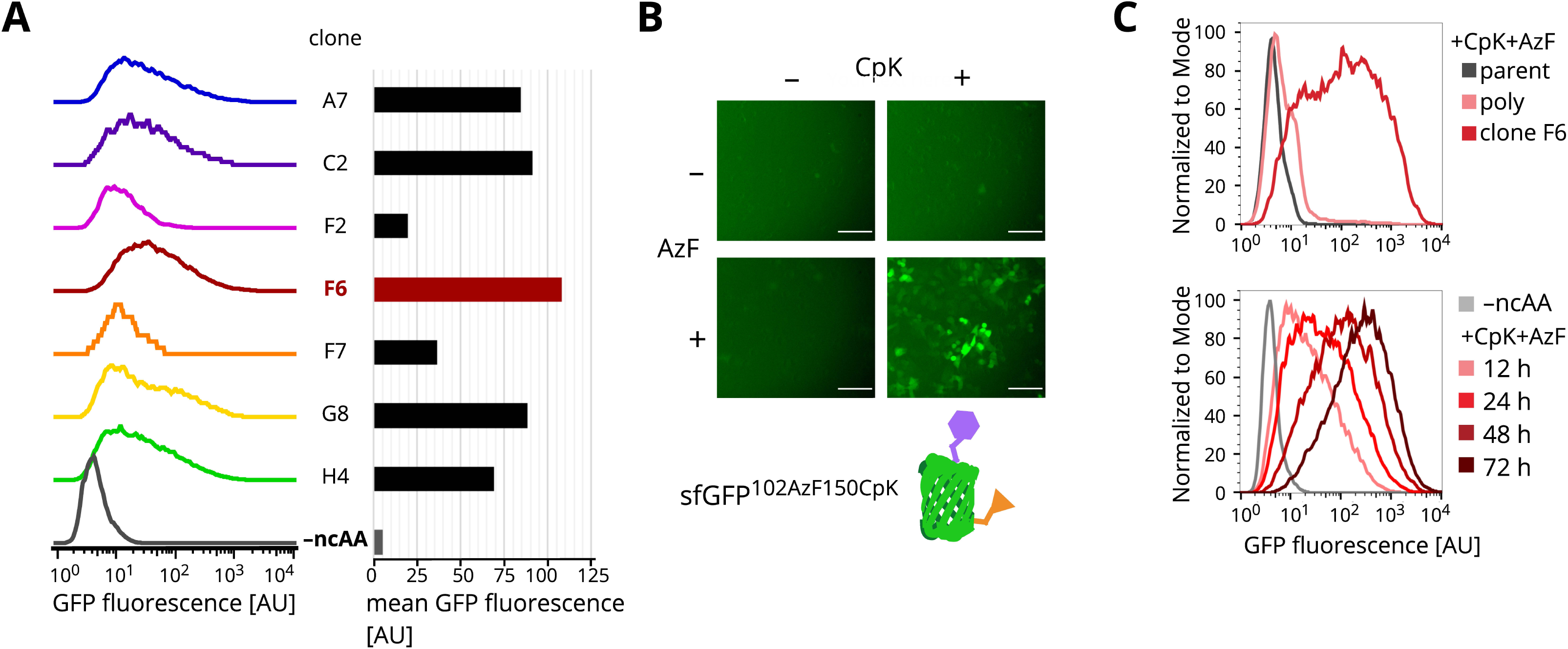
Dual suppression clonal cell line for GFP^102AzF150CpK^ expression in HCT116 cells. Related to Figure 4. A. Comparison of GFP fluorescence generated by dual amber and ochre suppression in the dual suppression clones sorted from a common polyclonal pool of tRNA^Pyl^_UUA_/PylRS ochre suppressor, tRNA^Tyr^_CUA_/AzFRS amber suppressor pair and GFP^102TAG150TAA^ reporter integrants generated by PB transgenesis. The clonal populations were grown in presence of 0.2 mM CpK and 0.5 mM AzF for 48 h. B. Fluorescence microscopy images of dual suppressor clonal F6 population grown without ncAA supplement, with either CpK or AzF or both for 48 h. White scale bars indicate 250 μm. C. Comparison of GFP fluorescence generated by dual amber and ochre suppression in the dual suppression cell polyclonal pool of tRNA^Pyl^_UUA_/PylRS ochre suppressor, tRNA^Tyr^_CUA_/AzFRS amber suppressor pair and GFP^102TAG150TAA^ reporter, the derived F6 clonal population or HCT116 parental cells. The populations were grown in presence of 0.2 mM CpK and 0.5 mM AzF for 48 h before measuring GFP fluorescence by flow cytometry. D. Time course of ncAA incorporation by the dual suppressor F6 clonal population, cells were grown without ncAAs or with 0.2 mM CpK and 0.5 mM AzF for 12 h, 24 h, 48 h or 72 h prior to measuring GFP fluorescence by flow cytometry.

**Figure S5.**
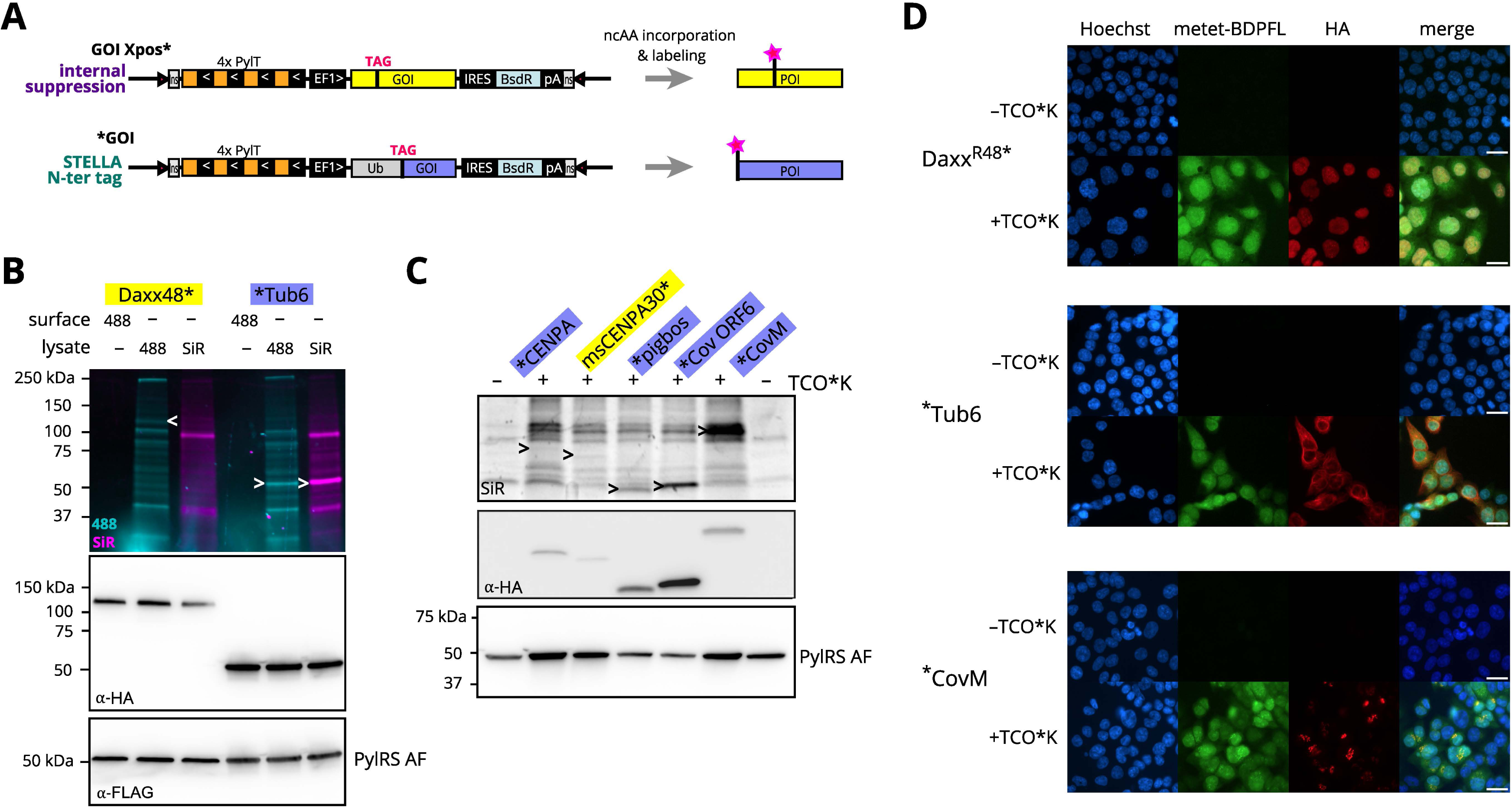
Amber suppression cell lines for bioorthogonal labeling of intracellular proteins. Related to Figure 5. A. Scheme illustrating construct design and ncAA labeling approaches for internal TAG and STELLA amber suppression. An asterisk (*) is used to indicate position of the amber codon and ncAA incorporation site. The amber stop codon signaling for ncAA incorporation can either be placed within the coding sequence of a gene of interest (GOI), producing the protein of interest (POI) in an ncAA-dependent manner, alternatively the POI can be expressed with an N-terminal ubiquitin-tag separated by an amber codon, leading to POI with ncAA at the N-terminus after ubiquitin processing (STELLA N-terminal tag) ^4^. B. Amber suppression PB integrant cell lines with tRNA^Pyl^/PylRS AF of Daxx^48TAG^-HA (Daxx48*) or STELLA N-terminally tagged Tub6-HA (*Tub6). Stable integrant populations recovered after 7 d selection were grown in presence of 0.1 mM TCO*K for 24 h and SPIEDAC-labeled on live cells with 4 μM tet488 or in lysate with 1 μM tet488 or 1 μM tetSiR. Lysate aliquots were separated by SDS-PAGE and fluorescence captured in-gel prior to membrane transfer and immunoblotting for the C-terminal HA-tag and the FLAG-PylRS AF. C. In-gel fluorescence and immunoblotting for further PB integrant amber suppression cells lines generated in HCT116 cells with tRNA^Pyl^/PylRS AF and STELLA N-terminally tagged CENPA, pigbos, *Cov*ORF6 and *Cov*M or murine CENPA Q30TAG (msCENPA30*). Stable integrant populations recovered after 7 d selection were grown in presence of 0.1 mM TCO*K for 24 h and SPIEDAC-labeled in lysate with 1 μM tetSiR. Lysate aliquots were separated by SDS-PAGE and in-gel fluorescence visualized prior to membrane transfer and western blot for the C-terminal HA-tag and the FLAG-PylRS AF. D. Fluorescence microscopy of amber suppressor HCT116 polyclonal pools encoding tRNA^Pyl^/PylRS AF and a protein of interest: Daxx^48TAG^ or STELLA N-terminally tagged Tub6 or *Cov*M. Cells were grown in presence of 0.1 mM TCO*K for 48 h. Cells were fixed with 4% PFA, permeabilized with 0.1% triton, blocked with 2% BSA and stained with 0.5 μM metetBDPFL, primary anti-HA and secondary 555-coupled antibodies. Prior to imaging on a Nikon Ti2, nuclei were counterstained with Hoechst 33342. White scale bars indicate 25 μm.

**Figure S6.**
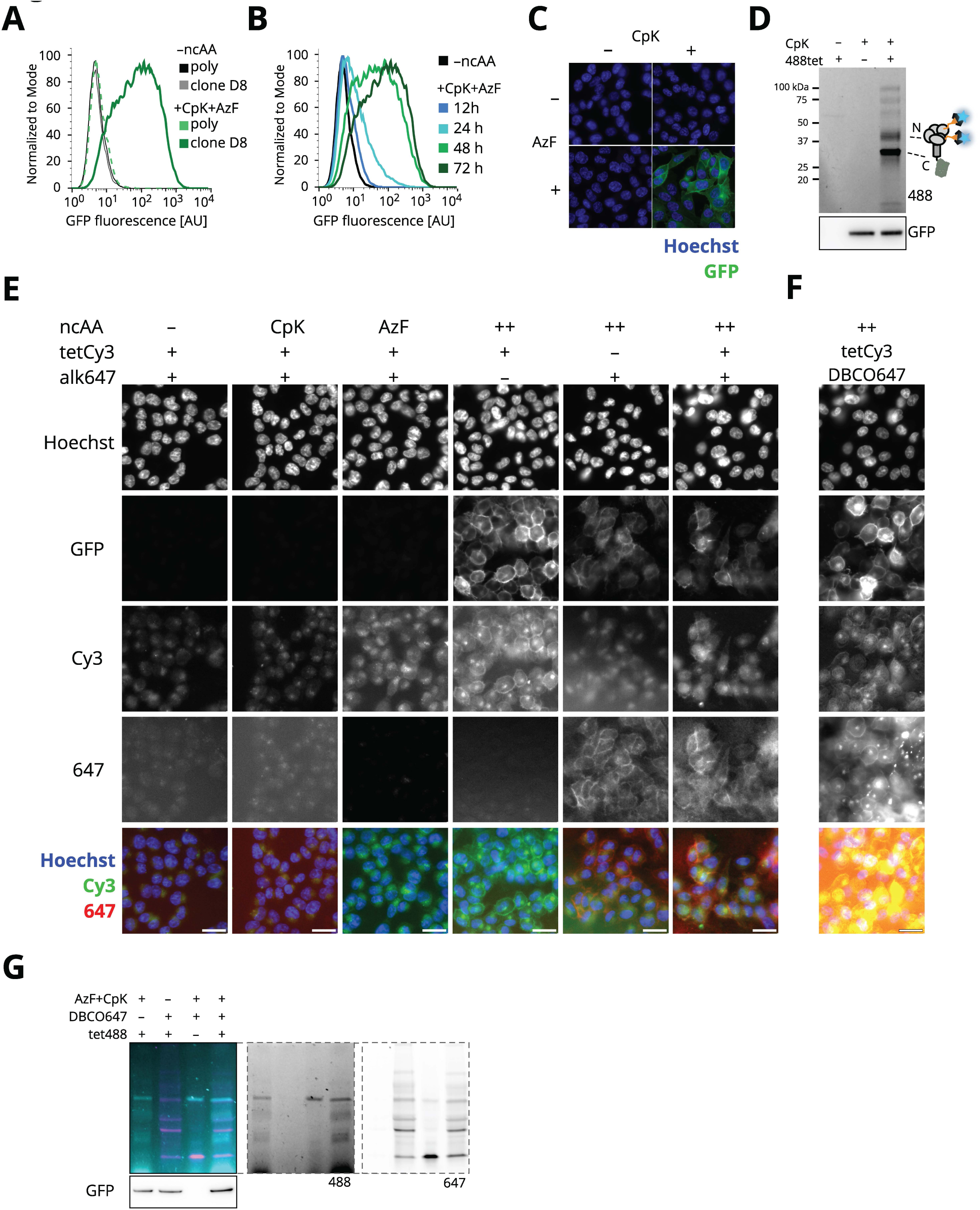
Dual suppression HCT116 clonal cell line for SynNotch^204CpK442AzF^ expression and live-cell dual bioorthogonal labeling. Related to Figure 6. A. Comparison of GFP fluorescence generated by dual amber and ochre suppression in the dual suppression cell polyclonal pool of M15_UUA_/PylRS ochre suppressor, tRNA^Tyr^_CUA_/AzFRS amber suppressor pair and SynNotch^204TAG442TAA^-GFP, the derived D8 clonal population or HCT116 parental cells. The populations were grown in presence of 0.2 mM CpK and 0.5 mM AzF for 72 h before measuring GFP fluorescence by flow cytometry. B. Time course of ncAA incorporation by the dual suppressor D8 clonal population, cells were grown without ncAAs or with 0.2 mM CpK and 0.5 mM AzF for 12 h, 24 h, 48 h or 72 h prior to measuring GFP fluorescence by flow cytometry. C. Fluorescence microscopy images of dual suppressor SynNotch^204TAG442TAA^-GFP clonal D8 population grown without ncAA, with either CpK or AzF or both ncAAs for 48 h. GFP fluorescene (green) and Hoechst 33342 nuclear staining (blue) are shown. D. Transient transfection of HEK293T cells with tRNA^Pyl^/PylRS and M15_UUA_/SynNotch^204TAA442TAG^-GFP plasmids (at 1:4 ratio) grown with and without 0.2 mM CpK. Live cells were surface labeled with 2 μM tet488 48 h post transfection. Aliquots of cell lysates were separated by SDS-PAGE, exposed at 460 nm (488) for in-gel fluorescence and immunostained for GFP and FLAG-RS after membrane transfer. E. Live-cell imaging of SPIEDAC-and CuAAC-labeled HCT116-derived dual suppressor clonal population with coding sequences for M15_UUA_/PylRS, tRNA^Tyr^_CUA_/AzFRS, and M15_UUA_/SynNotch^204TAA442TAG^-GFP integrated by PB transposition. Cells were grown for 72 h in the absence (–ncAA) or the presence of 0.2 mM CpK and 0.5 mM AzF. Surface labeling by SPIEDAC with 2 μM tetCy3 and CuAAC with 2 μM alk647 was carried out prior to Hoechst 33342 counterstaining, fixation with PFA and imaging. White scale bars indicate 25 μm. F. Conditions like E) but instead of CuAAC with alk647, AzF was labeled by SPAAC with 2 μM DBCO-647. White scale bars indicate 25 μm. G. SPIEDAC and SPAAC labeling on live HCT116 clonal SynNotch^204CpK442AzF^-GFP cells. grown in presence of 0.2 mM CpK and 0.5 mM AzF for 72 h. Cells were labeled with 2 μM tet488 and 2 µM DBCO-647. Lysates were separated by SDS-PAGE, in-gel fluorescence at 460 nm (488) and 630 nm (647) and immunoblotting against the C-terminal GFP are shown.

## Notes

### Competing Interest Statement

The authors have declared no competing interest.

### Summary of Updates

Corrected and updated text and figures. Added supplementary data.

